# Systematic drug profiling across BAF complex perturbations reveals distinct dependencies

**DOI:** 10.64898/2026.03.11.710932

**Authors:** Katharina Spang, Caroline Barry, Christina Ntasiou, Eleni Aretaki, Mara Wolf, Eric Schumbera, Fridolin Kielisch, Lena Bonn, Mareen Welzel, Frank Rühle, Christian Schäfer, Katja Luck, Sandra Schick

## Abstract

Chromatin remodeling by BAF complexes regulates access to DNA with implications in transcription, replication and the sensing and repair of DNA lesions, processes that are closely coupled to cell growth control. BAF complexes exist in three major subtypes each composed of a unique subset of about a dozen subunits. While for individual BAF subunits direct functions in DNA repair and cell growth have been described, we lack a more systematic interrogation of BAF perturbation and its implications in these processes. To fill this need, we subjected an isogenic BAF knockout collection to treatment with diverse genotoxic and cytotoxic compounds quantifying proliferation, cell cycle phase distributions, cell fate decisions, DNA double-strand break signaling, and transcriptional responses. The screen uncovered vulnerabilities of selected BAF-deficient cells to MEK and EGFR pathway inhibition, as well as unexpected resistance toward checkpoint inhibitors and certain DNA-damaging agents. Notably, loss of distinct ARID paralogs produced markedly divergent phenotypes, indicating limited functional redundancy and subtype-specific contributions to genome stability and cell cycle regulation. Collectively, these results provide a comparative resource linking BAF complex composition to genotoxic and cytotoxic stress responses, highlighting candidate synthetic-lethal interactions and generating hypotheses for mechanistic studies of BAF complexes in genome maintenance and cell growth control.

## Introduction

Numerous processes act on the genome to regulate access to DNA, DNA repair, transcription, and replication ensuring genome integrity, which is essential for physiological cellular function and correct inheritance of genetic material. Genome integrity processes are, thus, closely integrated with checkpoint signaling where unrepaired DNA damage causes cell cycle arrest, senescence, and apoptosis. Genome instability in combination with unregulated cell growth are hallmarks of cancer^1^. DNA accessibility is a major contributing but underexplored factor to genome integrity. The level of chromatin compaction and nucleosome composition strongly influences transcription and replication initiation as well as their progression, and regulates access to DNA lesions for repair^2^.

The human ‘BRG1- or BRM-associated factor’ (BAF) complexes, also known as ‘switch/sucrose-non-fermenting’ (SWI/SNF) complexes, are prominent chromatin remodeling complexes that use ATP hydrolysis to slide or evict nucleosomes from the DNA^3^. In total, 29 subunits have been identified to-date that can assemble into three different BAF complex subtypes^3^. Each BAF complex consists of an ATPase module, an ARP module, and a core module. The ATPase module is composed of SMARCA4 or its paralog SMARCA2. Both subunits are common to all BAF subtypes and carry the ATP hydrolysis activity^4^. They interact with the nucleosome and connect to all other modules and thus, can be seen as the backbone of BAF complexes^5,6^. The ARP module consists of ACTL6A/B, ACTB, and BCL7A/B/C regulating the ATPase activity. The core module provides the structural foundation of BAF complexes and assists in nucleosome binding, DNA targeting and regulation of BAF complex function via physical interactions with DNA, histone marks, transcription factors, and other regulatory proteins or RNA^3,5,6^. The subunit composition of the core module defines three BAF subtypes: ARID1A/B and DPF1/2/3 are exclusively incorporated into the canonical BAF (cBAF), ARID2, BRD7, PHF10, and PBRM1 define the polybromo-associated BAF (PBAF), and BICRA/L and BRD9 form the non-canonical BAF (ncBAF)^7,8^. Selected incorporation of paralogous subunits into the BAF subtypes varies between cell types and developmental stages. For instance, a switch from ACTL6A to ACTL6B incorporation into cBAF was observed during differentiation from neural progenitors to neurons^9^. However, how different subunit composition drives differential function of BAF complexes is largely unresolved.

Given tight connections between chromatin remodeling, genome integrity, and cell growth, it is not surprising that mutations in BAF complex subunits are associated with tumorigenesis. In fact, more than 20% of all cancers carry at least one mutation in a BAF subunit, with ARID1A being most frequently mutated^10^. Mutation frequency of individual BAF subunits strongly varies by cancer type. While SMARCA4 is mutated in 90–100% of small cell ovarian cancers^11^, SMARCB1 is mutated in nearly all malignant rhabdoid tumors for which its loss was demonstrated to be the key driver^12,13^. Mechanistically, BAF perturbation is hypothesized to alter gene expression programs driving the deregulation of cell growth factors thereby contributing to uncontrolled cell proliferation. For instance, ARID2 downregulation was found to accelerate G1/S transitions via transcriptional upregulation of Cyclin D1, E1, and CDK4^14^. ARID1A, but not ARID1B, has been implicated in enforcing G1 arrest through induction of p21 and repression of E2F target genes^15^. Finally, ARID1A loss was also found to increase EGF-mediated growth signaling^16^. However, the direct involvement of individual BAF subunits, notably SMARCA4 and ARID1A, in DNA damage sensing and repair processes also suggests that an increase in genome instability upon BAF loss contributes to carcinogenesis. For example, SMARCA4 loss enhanced or reduced γH2AX levels in response to DNA double strand break (DSB) induction depending on cellular context and time after damage^17,18^. Furthermore, phosphorylation of SMARCA4 by ATM was found to promote SMARCA4 recruitment to DSBs^19^. On the other hand, ARID1A is recruited to DSBs via interaction with ATR and loss of ARID1A resulted in defective homologous recombination-mediated repair (HR)^20^. While these focused studies significantly improved our understanding of BAF function beyond chromatin remodeling, we still lack a more systematic interrogation of BAF perturbation and its implications on genome stability and cell growth to better understand which BAF subunits and subtypes directly function in these processes and how.

Improved understanding of these crosstalks between BAF and genome maintenance as well as with cell growth regulation has high therapeutic potential through the exploration of synthetic lethalities. For example, treatment with PARP and ATR inhibitors led to sensitivities specifically in a BAF-perturbed genetic background^20,21^. Likely more synthetic lethal relationships between BAF loss and DNA repair or cell growth signaling pathways exist yet remain to be explored. To advance our understanding of BAF subunit and subtype-specific roles in these processes, we set out to screen a comprehensive BAF knockout (KO) collection in an isogenic background against various genotoxic and cytotoxic compounds quantifying implications on cell growth, cell cycle phases, cell fate, DSB signaling, and transcription. We identified sensitivities of specific BAF subunit loss upon treatment with MEK and EGFR inhibitors and discovered unexpected resistances in combination with cell cycle checkpoint inhibitors and selected genotoxic agents. Most notably, we describe highly differential responses to these treatments depending on the type of ARID subunit loss pointing to incomplete compensatory mechanisms of these paralogs and unique roles in genome stability and cell cycle regulation. This resource enabled the formulation of various hypotheses that will form the basis for future studies to mechanistically disentangle the function of distinct BAF complexes beyond chromatin remodeling.

## Results

### Profiling of cell physiologic and transcriptomic changes upon BAF subunit loss

To systematically investigate the impact of individual BAF subunit perturbation on cell physiology including cell growth, cell cycling, cell fate, and genome stability, we used an existing CRISPR/Cas9-generated BAF subunit KO collection in the haploid leukemia cell line HAP1^8^. We selected the cBAF-specific subunits ARID1A and ARID1B, the PBAF-specific subunits ARID2, BRD7, PHF10, and PBRM1, the ncBAF-specific subunit BRD9 as well as the common BAF subunits SMARCD1 (the only SMARCD subunit that is incorporated in ncBAF), SMARCC1, and SMARCA4 to profile effects of BAF subtype-specific as well as of common BAF subunit perturbations (Fig. 1a). Since HAP1 cells can spontaneously turn diploid, we cultured and FACS-sorted all selected HAP1 BAF KO cell lines to obtain diploid cell lines that are more stable in their ploidy and reflect the ploidy of most human somatic cells (Fig. S1). To investigate the impact of individual BAF subunit KO on cell proliferation, we monitored increases in confluency over five days observing that compared to WT, ARID1B KO cells displayed a significantly increased growth rate, while SMARCC1 cells displayed a significantly decreased growth rate and max capacity. SMARCA4 KO cells displayed significantly increased max capacity, but no alterations in growth rate (Fig. 1b, Fig. S2a, Table S1). Alterations in the growth behavior of these three BAF KO cell lines are further characterized by an increase in the number of nuclei and decrease in nucleus area for ARID1B KO cells, the inverse for SMARCC1 and SMARCA4 KO cells based on immunofluorescence microscopy analysis (Fig. S2b-c, Table S1). KO of all other tested BAF subunits did not show significant changes in growth rate, suggesting that without additional cellular stresses perturbation of most BAF subunits in the HAP1 context has little to no immediate impact on cell growth (Fig. 1b).

**Figure 1:**
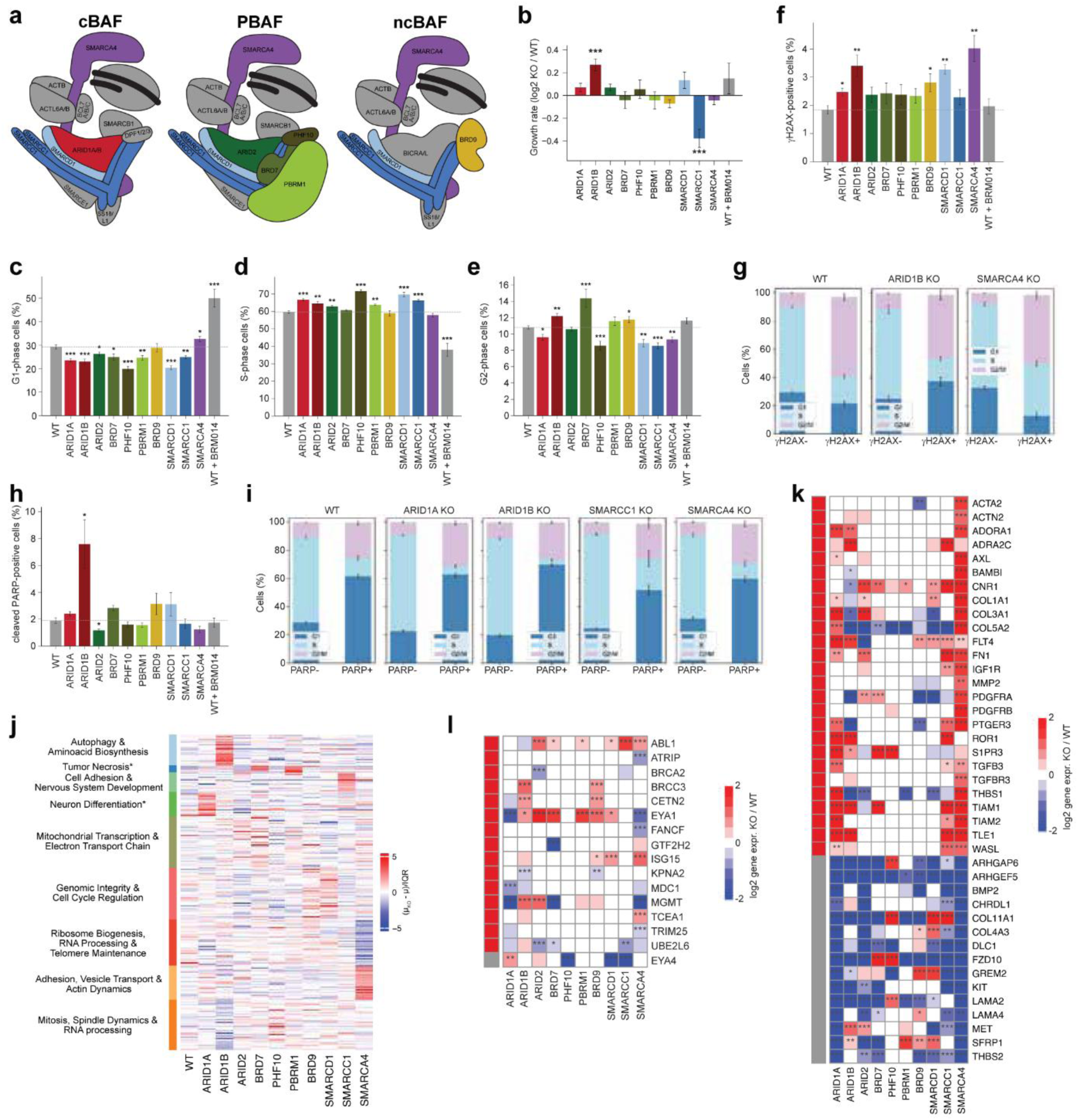
Phenotyping of a HAP1 BAF KO cell line panel. **a.** Schematic showing the three BAF subtypes with subunits highlighted in color that are part of the HAP1 KO collection. **b.** Bar plot showing log2 fold changes of mean growth rates of BAF KO vs WT cells estimated from confluency curves obtained from live-cell imaging. * p-value < 0.05, ** p-value < 0.01, *** p-value < 0.001. Error bars indicate STE, n ≤ 3. **c-e.** Bar plots visualizing mean fraction of cells in G1, S, and G2 cell cycle phase in HAP1 WT and BAF KOs. Measurements obtained from flow cytometry quantifying EdU incorporation and Hoechst stain for DNA content. Significance determined using t-test comparing KO vs WT, p-values as in **b**. Error bars indicate STE, n≥3. **f.** Bar plot showing mean fraction of γH2AX-positive cells upon BAF KO and WT. Measurements obtained from flow cytometry stratifying cells by stains for γH2AX. Significance determined using t-test comparing KO vs WT, p-values as in **b**. Error bars indicate STE, n≥3. **g.** Bar plots showing the fractions of cells in G1, S, and G2 phase for WT (left), ARID1B KO (middle), SMARCA4 KO (right) cells. Cells are split into γH2AX negative or positive cells upon treatment with DMSO. Error bars indicate STE, n≥3. **h.** Bar plot visualizing mean fraction of cleaved PARP-positive cells upon BAF KO and WT. Measurements obtained from flow cytometry stratifying cells by stains for cleaved PARP. Statistics as in **f.** Error bars indicate STE, n≥3. **i.** Bar plots visualizing the fractions of cells in G1, S, and G2 phase for WT, ARID1A KO, ARID1B KO, SMARCC1 KO, and SMARCA4 KO (from left to right) cells. Cells are split into PARP negative and positive cells upon treatment with DMSO. Error bars indicate STE, n≥3. **j.** Heatmap showing Preferential Expression Score (PES) upon BAF KO for protein-coding genes with |PES| > 1 in at least one cell line. GO term enrichment analysis was performed for each cluster and representative GO terms for each cluster are indicated. * indicates GO terms that were not significant. **k-l.** Heatmaps containing log2 fold changes in gene expression upon BAF KO compared to WT shown for selected subsets of differentially expressed genes based on Reactome pathway annotations (cell growth signaling in **k**, DNA repair in **l**) clustered into 2 groups. Significance determined from DESeq2, p-values as in **b**, n = 3 (PBRM1 KO n=2).

Cell cycle analysis of all BAF KO cell lines using flow cytometry showed for most BAF KOs a significantly higher fraction of cells in S phase compared to WT, and a reduced fraction of cells in G1 and to some extent also in G2 (Fig. 1c-e, Fig. S2d, Table S1). Since the growth rate was not significantly increased for most BAF KO cells compared to WT, this suggests that most BAF subunit KOs lead to a prolonged S and shorter G1 and G2 phases. All BAF KO cell lines also displayed elevated levels of γH2AX in flow cytometric analysis although significant only for ARID1A/B, BRD9, SMARCD1, and SMARCA4 KO cells (Fig. 1f). This suggests that prolonged S phase can be a result of slowed down replication due to increased levels of DSBs. We furthermore observed that for WT and ARID1B KO cells, γH2AX positive cells accumulate the most in G2 and less in replicating cells. In comparison to that, the percentage of γH2AX positive cells is higher in SMARCA4 KO replicating cells, suggesting that DSBs in SMARCA4 KO cells trigger less G2/M and G1 checkpoints leading to more cells entering S phase with DSBs or that DSBs occur more frequently in S phase (Fig. 1g). Interestingly, the fraction of S phase cells in SMARCA4 KO is not altered, yet nucleus area and max confluency are increased (Fig. 1d, Fig. S2a,c). ARID1B KO cells displayed a significant increase in apoptotic cells as determined by staining for cleaved PARP in flow cytometric analysis, while ARID2 KO cells displayed slightly reduced apoptotic levels compared to WT (Fig. 1h, Table S1).

We furthermore observed that cleaved PARP-positive ARID1B KO cells accumulate in G1 at a higher percentage compared to WT with almost no cells in S phase that undergo apoptosis (Fig. 1i). We made similar observations for ARID1A KO cells but not for any of the other BAF KO cell lines (Fig. 1i), suggesting that ARID1A and ARID1B KO lead to altered induction of apoptosis. Interestingly, specific inhibition of the ATPase activity of SMARCA4/2 by the small molecule BRM014^22,23^ showed a significant reduction of cells in S phase and increase of cells in G1 but no changes to γH2AX and cleaved PARP levels (Fig. 1c-d, f, h). Furthermore, growth rate and max capacity of HAP1 cells treated with BRM014 were not significantly different compared to untreated HAP1 cells (Fig. 1b, Fig. S2a). This suggests that perturbation of BAF by subunit KO leads to cell cycle, growth, and genome stability alterations that are different from selective perturbation of the ATPase-driven nucleosome remodeling activity.

We reanalyzed an existing RNA-seq dataset of the HAP1 BAF KO collection^8^ to assess differences in gene expression changes upon loss of individual BAF subunits. As previously reported, we observed the most differences in gene expression compared to WT upon loss of SMARCA4, the three ARID subunits, and SMARCC1 (Fig. S2e, Table S2). The smallest number of differentially expressed genes was observed for the PBAF-specific subunits PBRM1 and PHF10 (Fig. S2e). In line with this, we also observe that the transcriptome profile of SMARCA4 KO cells is the most different to the other BAF KO cell lines, while PBAF-specific KO cell lines (ARID2, PBRM1, BRD7) cluster together (Fig. S2f). ARID1B, ARID1A, and SMARCC1 KO also lead to distinct transcriptome profiles that do not cluster with any other BAF KO cell line. Interestingly, ARID1B KO gene expression is the most different to SMARCA4 KO cells while ARID1A KO cells are the most similar to SMARCA4 KO cells among all profiled BAF KOs (Fig. S2f). Significantly upregulated genes upon BAF KO were enriched for neuronal system and muscle tissue development including functional terms such as axogenesis, neural crest differentiation and cell adhesion functions. Genes functioning in the negative regulation of focal adhesions or in cognitive functions were significantly downregulated (Fig. S2g). To further identify genes that are uniquely differentially up- or downregulated upon individual BAF subunit KO, we determined how many interquartile ranges this gene is more or less expressed in this condition compared to its average expression in all other conditions^24^. Clustering all genes with a preferential expression score of >1 or <1 revealed specific upregulation of autophagy-related genes and downregulation of mitosis and spindle dynamics-related genes in the ARID1B KO background (Fig. 1j, Table S3). Functional terms relating to telomere maintenance, ribosome biogenesis and RNA processing were specifically downregulated in the SMARCA4 KO background (Fig. 1j).

Due to observed alterations in cell growth and cycling of various BAF KOs, we more specifically investigated differential expression of genes functioning in cell growth signaling pathways. We observe downregulation of Wnt and BMP signaling pathway members (FZD10, SFRP1) for most BAF KO cell lines as well as downregulation of extracellular matrix and cell adhesion factors (Fig. 1k). A set of genes that shows specific upregulation for ARID1A, SMARCC1 and most strongly for SMARCA4 KO cells functions in growth factor and receptor tyrosine signaling (PDGFRB, FLT4, AXL, TFGBR3, IGF1R), extracellular matrix and cytoskeleton remodeling as well as G protein-coupled receptor (GPCR) signaling, suggesting that loss of core BAF subunits requires substantial remodeling of growth signaling pathways for cells to survive. We also investigated differential expression of genes functioning in DNA damage repair. In comparison to all protein coding genes and genes functioning in cell growth signaling, where SMARCA4, SMARCC1, ARID1A and ARID1B KO cells showed the most differentially expressed genes, here, ARID1B and SMARCA4 KO cells displayed the highest number of differentially expressed genes functioning in DNA damage repair (Fig. S2h). In particular, we observed a marked downregulation of the O6-methylguanine-DNA methyltransferase (MGMT) in the ARID1A, BRD7, and SMARCA4 KO background (Fig. 1l). MGMT repairs O6-methylguanines, which otherwise would cause mismatches in the DNA upon replication^25^, suggesting that SMARCA4 and ARID1A-deficient HAP1 cells are more sensitive to O6-methyl-alkylating agents such as MNNG. We also observed differential expression of the Tyrosine phosphatase EYA1 and EYA4 in various BAF KO cell lines (Fig. 1l). EYA1/4 are known to dephosphorylate H2AX at Y142 enabling efficient recruitment of downstream DNA repair factors by γH2AX signaling^26^. Conversely, persistent phosphorylation of Y142 upon DNA damage triggers apoptotic signals potentially deregulating apoptotic signaling in BAF KO cells upon genotoxic stress. In summary, we observed BAF subunit loss-specific alterations in cell growth, cell cycling, γH2AX levels, and apoptosis. While no BAF subtype-specific trends were observed, we found that perturbation of SMARCA4, SMARCC1, and the ARID subunits led to the most significant and often different alterations in cell physiology, while perturbation of other PBAF and ncBAF-specific subunits for these readouts rarely resulted in significant phenotypes.

### Identification of vulnerabilities upon BAF loss in published drug screening data

To further identify BAF subunit loss-specific alterations on cell physiology we aimed to identify sensitivities and resistances in a BAF perturbation background upon treatment of cells with genotoxic or cytotoxic compounds. To this end, we analyzed the PRISM dataset in which alterations in cell growth were quantified for 919 cancer cell lines in response to treatment with 1,448 approved cancer drugs or drugs in clinical trials^27^. We identified significant differences in the sensitivity to treatment with 24 compounds between cell lines of the top and bottom 20th percentile when ranked by gene expression for a given BAF subunit of our panel. We found that reduced expression of the three ARID subunits and especially of ARID2 correlated with an increased sensitivity to MEK inhibitors (Fig. 2a). Interestingly, cells with reduced expression of the PBAF-specific subunit PBRM1 displayed sensitivity when treated with EGFR or VEGFR inhibitors (Fig. 2a). Furthermore, we observed that genotoxic agents such as the DNA alkylating agent Melphalan leading to DNA interstrand crosslinks, the PARP inhibitor Niraparib, as well as the Topoisomerase 2 (Top2) inhibitor Voreloxin led to improved cell growth upon reduced BAF subunit expression. This was significant for SMARCC1 and SMARCA4 (Fig. 2a). Next, we analyzed results from a genome-wide CRISPR KO drug sensitivity screen conducted in RPE-1 cells against 27 genotoxic agents (hereafter referred to as the Olivieri dataset)^28^. Out of the 27 tested compounds 6 showed significant sensitivity upon KO of at least one of the BAF subunits in our panel. Interestingly, none resulted in significant resistances using reported cutoffs, however, treatment of cells with the Topoisomerase 1 (Top1) inhibitor Camptothecin in PBRM1 KO cells and the DNA-alkylating agent IlludinS in ARID1A KO cells resulted in resistances close to the significance cutoff (Fig. 2b). Interestingly, treatment of cells with IlludinS and Camptothecin resulted in sensitivities specific to perturbation of the ncBAF-specific subunit BRD9 and SMARCD1, while treatment with the NEDD8-activating enzyme inhibitor MLN4924 (MLN) resulted in sensitivities upon individual loss of all four PBAF-specific subunits (Fig. 2b).

**Figure 2:**
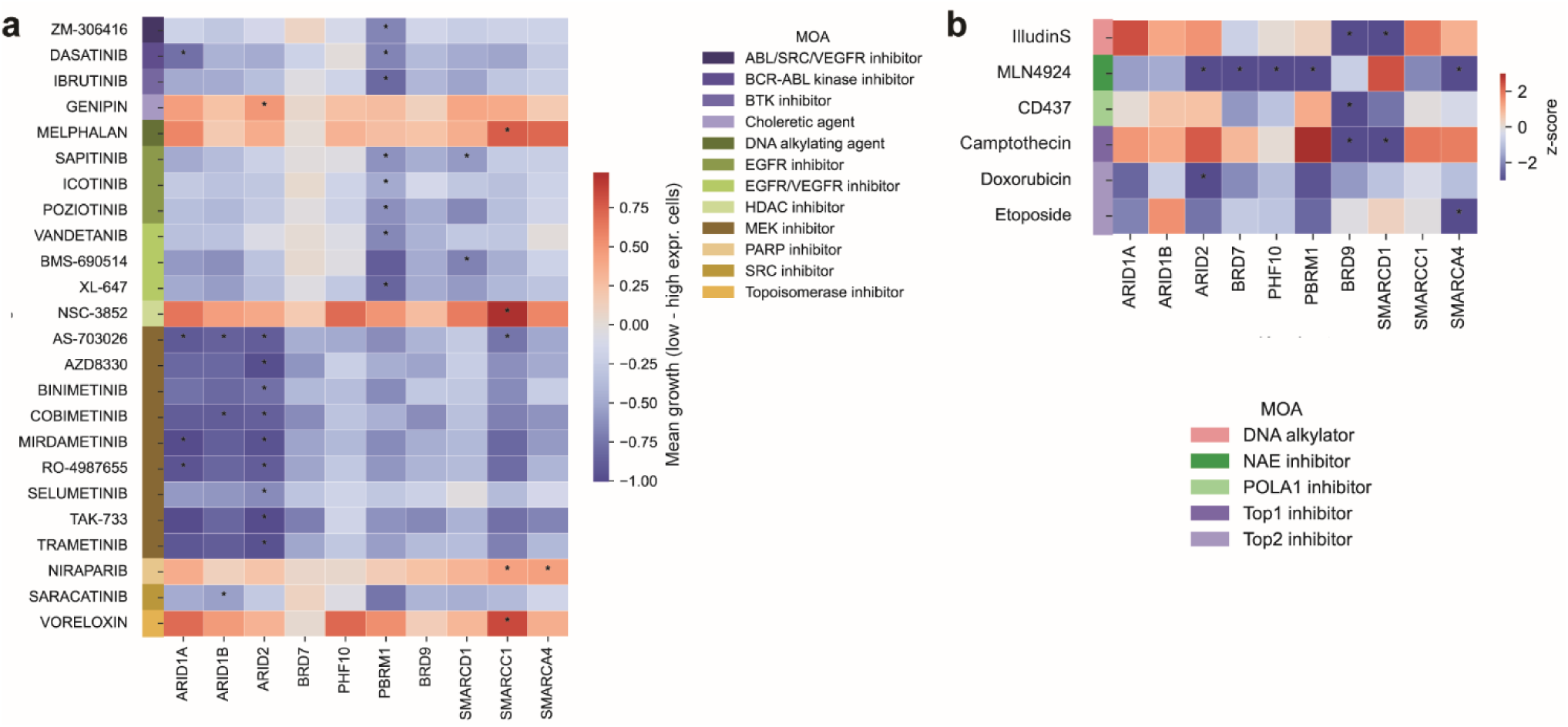
Compound sensitivity upon BAF perturbation across public datasets. **a.** Heatmap showing mean difference in growth response of cells treated with various compounds subtracting growth of cells with high BAF subunit gene expression from those with low levels across an array of cancer cell lines from the PRISM dataset. All significant compounds across the BAF subunit panel are shown. Significance computed with Mann–Whitney U-test, p-values corrected for multiple testing, * p-value < 0.01. **b.** Heatmap visualizing z-scores obtained from Olivieri dataset for significant compounds across the BAF KO panel. Z-scores < -3 or > 6 were deemed significantly sensitizing or resisting hits, respectively, according to Olivieri et al, FDR < 15%.

### Profiling of cell growth alterations upon BAF subunit loss and drug treatment

To confirm previously observed and identify novel vulnerabilities of BAF subunit loss upon treatment with cytotoxic or genotoxic agents in an isogenic background, we selected 21 compounds that impact cell cycle regulation, growth signaling, or genome stability (Fig. 3a). These compounds were selected based on results from the PRISM and Olivieri dataset analysis as well as from previously reported implications of individual BAF subunits in the repair of DSBs and genome maintenance at telomeres^29^. To systematically quantify positive or negative effects of compound treatment on cell growth upon individual BAF subunit loss, we implemented a high-throughput (HTP) screening platform using the Incucyte live-cell imaging system (Fig. 3b). Cells were treated 24h after seeding with varying concentrations of a compound and confluency was quantified from brightfield microscopic images taken every 6 hours over a total time-course of 96h after treatment (Fig. 3b). In total, we performed 172,968 confluency measurements that we analyzed and integrated using a computational pipeline (GrowPipe) developed in R to quantify growth phenotypes. Clustering all conditions (21 compounds x 10 BAF KOs) by their growth effect compared to WT (and normalized to treatment with DMSO) across four time points and the ratio of 96h over 48h confluency post-treatment resulted in 9 clusters of compound-BAF combinations (Fig. 3c-d, Fig. S3, Fig. S4a, Table S4). The highest sensitivity in this screen was observed for SMARCC1 KO cells upon treatment with Cobimetinib forming cluster 1 (Fig. 3c,e). More than half of all conditions in cluster 2 consist of PBAF-specific subunits that displayed less growth compared to wild type (although not significant) upon treatment with MLN, consistent with observations from the Olivieri dataset, and upon treatment with Lapatinib or Sapitinib, two EGFR inhibitors, consistent with observations from the PRISM dataset (Fig. 3c). Cluster 3 is dominated by increasing sensitivity over time of SMARCC1 KO cells when treated with various different compounds (10/32 conditions). Nine out of all 32 conditions in this cluster also consist of MEK inhibitor treatment in combination with various different BAF subunit KOs such as SMARCA4, ARID1A and ARID2 consistent with observations from the PRISM dataset. Of note, while differences in growth were not significant for individual time points in this cluster, the ratio of confluency at the 96h over the 48h time point was significantly lower for treated KO versus treated WT cells (each normalized to DMSO) for most conditions in this cluster indicating worse growth at later time points compared to WT (Fig. 3c, e). Cluster 4 displays almost no significant changes in growth for any of the 61 conditions in this cluster, which are dominated by the PBAF-specific KOs BRD7 and PHF10 as well as the ncBAF-specific KOs BRD9 and SMARCD1 when treated with Top2 inhibitors, MLN, and the telomerase inhibitor BIBR1532. Clusters 5 to 9 are dominated by less growth impairment in the KOs, with cluster 5 displaying milder yet persistent resistance from the 48h to the 96h timepoint while clusters 6-9 displayed the strongest resistances (Fig. 3c). Resistances are observed for ARID1A and ARID1B but not ARID2 KO cells when treated with the cell cycle inhibitors Palbociclib and Paclitaxel (clusters 6 and 8, Fig. 3d,f). Of note, while ARID1B and ARID1A display accelerated recovery from Palbociclib treatment compared to wild type, ARID2 and PBRM1 instead show a delayed but strong growth defect (Fig. 3f). ARID1B, ARID2, and SMARCA4 but not ARID1A KO cells showed resistance upon treatment with the Top1 inhibitors Camptothecin and Topotecan (clusters 5 to 8) (Fig. 3c-d,g), while SMARCA4, SMARCC1, ARID1A, and ARID1B KO cells displayed strong resistance upon treatment with the PARP inhibitor Niraparib (clusters 8 and 9) (Fig. 3c-d,h). These results suggest that observed sensitivities for MEK and EGFR inhibitors in cancer cell lines persist in isogenic BAF KO cells and thus might be driven by BAF perturbation. At the same time, our results discover putative resistances to cell cycle checkpoint, PARP, and Top1 inhibition, especially upon KO of SMARCA4, SMARCC1, ARID1A and ARID1B.

**Figure 3:**
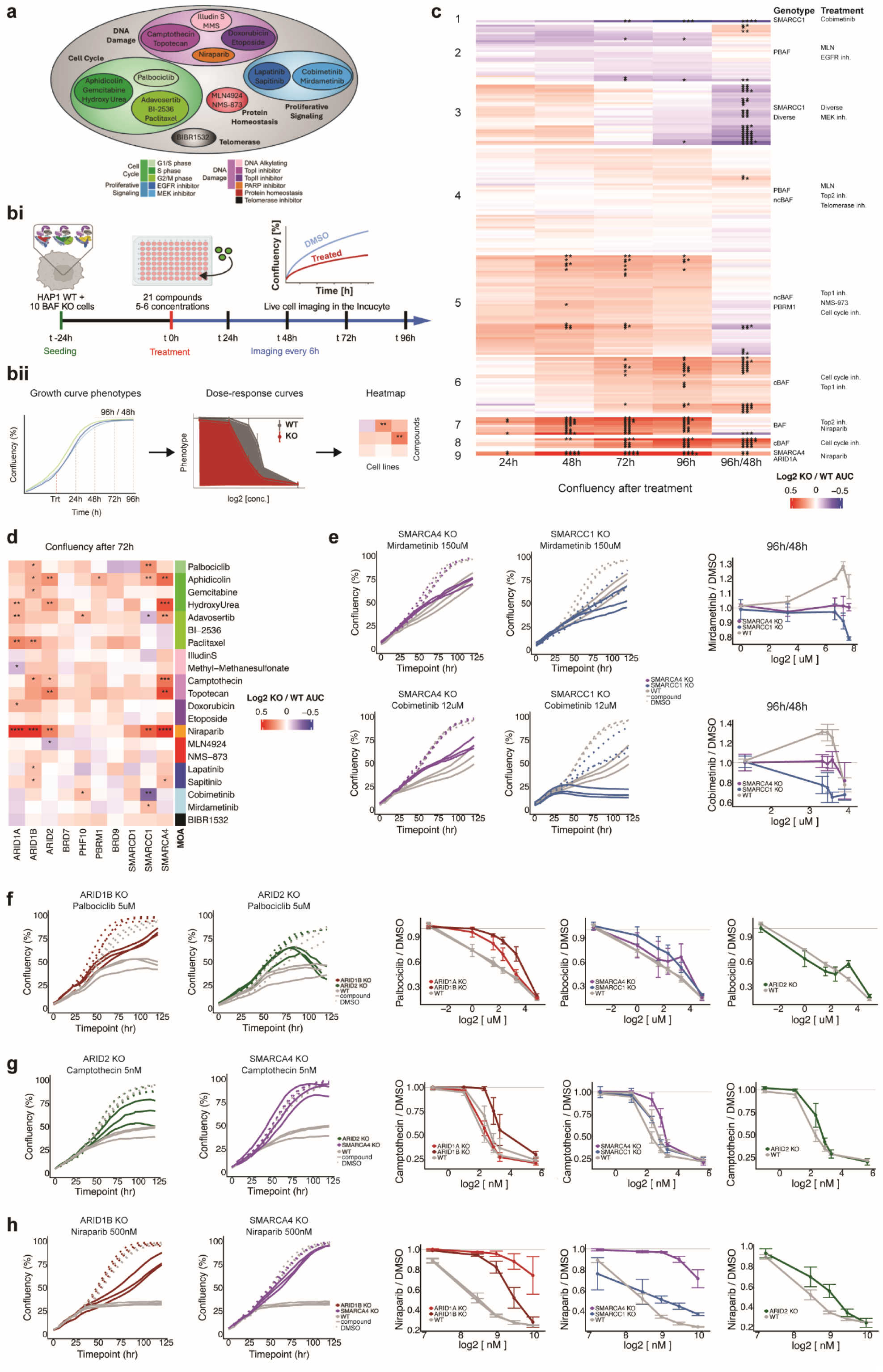
Alterations in cell growth upon BAF KO and treatment. **a.** Overview of the 21 compounds selected for sensitivity screens using cell growth as readout. **b.** Schematic showing the live-cell imaging screen setup (**i**) and computational pipeline to quantify cell growth (**ii**). **c.** Heatmap showing Area Under the dose response Curve (AUC) of confluences measured for BAF KO over WT for different treatments and time points after treatment, normalized each to DMSO and clustered into nine groups. Significance determined using t-test, p-values corrected for multiple testing, * p-value < 0.1, ** p-value < 0.05, *** p-value < 0.01 and **** p-value < 0.001, n=3, with exception for ARID2 KO & BRD9 KO treated with Etoposide and ARID1A KO treated with Niraparib where n=2. **d.** Heatmap showing growth effects as in **c** at 72h post-treatment. Statistics as in **c**. **e.** Growth curves of biological replicates and dose response curves for ratio of confluency after 96hr of treatment / confluency after 48hr of treatment for representative sensitization phenotypes (cluster 1 and 3). Error bars indicate STD, n=3. **f-h.** Growth curves of biological replicates and dose response curves for confluency after 96h of treatment for representative resistance phenotypes, clusters 6 and 8 (**f**), clusters 6 and 7 (**g**), clusters 8 and 9 (**h**). Error bars indicate STD, n=3. 3a) Created in BioRender. Bonn, L. (2026) https://BioRender.com/hwxgjzu 3bi) Created in BioRender. Schick, S. (2026) https://BioRender.com/nwbeuxg

### Phenotyping of alterations in cell cycle, DSB and apoptosis upon BAF perturbation and drug treatment

The previously described results indicate that the cellular dynamics of BAF KO cells in response to treatment with cell cycle, proliferative perturbants or DNA damaging agents substantially differ. While treatment with MEK and EGFR inhibitors resulted in sensitivities at later time points (i.e. 96 vs 48h, cluster 2), cell cycle inhibitors caused significant resistance starting at 72h post-treatment (cluster 5) and treatment with DNA damaging agents caused resistance as early as 48h after treatment (clusters 6 and 7). To further dissect early changes in cell fate signaling and DNA damage induction two days post-treatment and to understand how those might drive observed changes in cell growth, we conducted HTP flow cytometric analysis of selected BAF KO cells and compounds (Fig. 4a, S2d). We selected seven BAF subunits that resulted in significant growth alterations in the cell growth screen (these are SMARCA4, SMARCC1, the three ARID subunits, PBRM1, and BRD9) and chose one to two compounds per mode of action, ten in total. After treatment of cells for 48h and 1h EdU labeling, cells were stained for γH2AX to quantify DNA damage as a surrogate marker for DSB induction and activation of the DNA damage response, cleaved PARP to measure induction of apoptosis as well as EdU incorporation upon DNA synthesis to identify cells in S phase, and Hoechst stain to quantify DNA content and separate cells into either G1 or G2. Percentages of cells were determined for a given treated BAF KO cell line and compared to WT treated, both normalized to treatment with DMSO (Table S5). For conditions that were subjected to both, Incucyte and flow cytometry screens, we observed a strong negative correlation between increased fractions of cells undergoing apoptosis between BAF KO and WT upon treatment and increases in confluency (Fig. S4b). A significant positive correlation was observed for increases in fractions of cells in S phase between BAF KO and WT upon treatment and increases in confluency across conditions (Fig. S4c), supporting high levels of reproducibility of phenotypes measured across different platforms.

**Figure 4:**
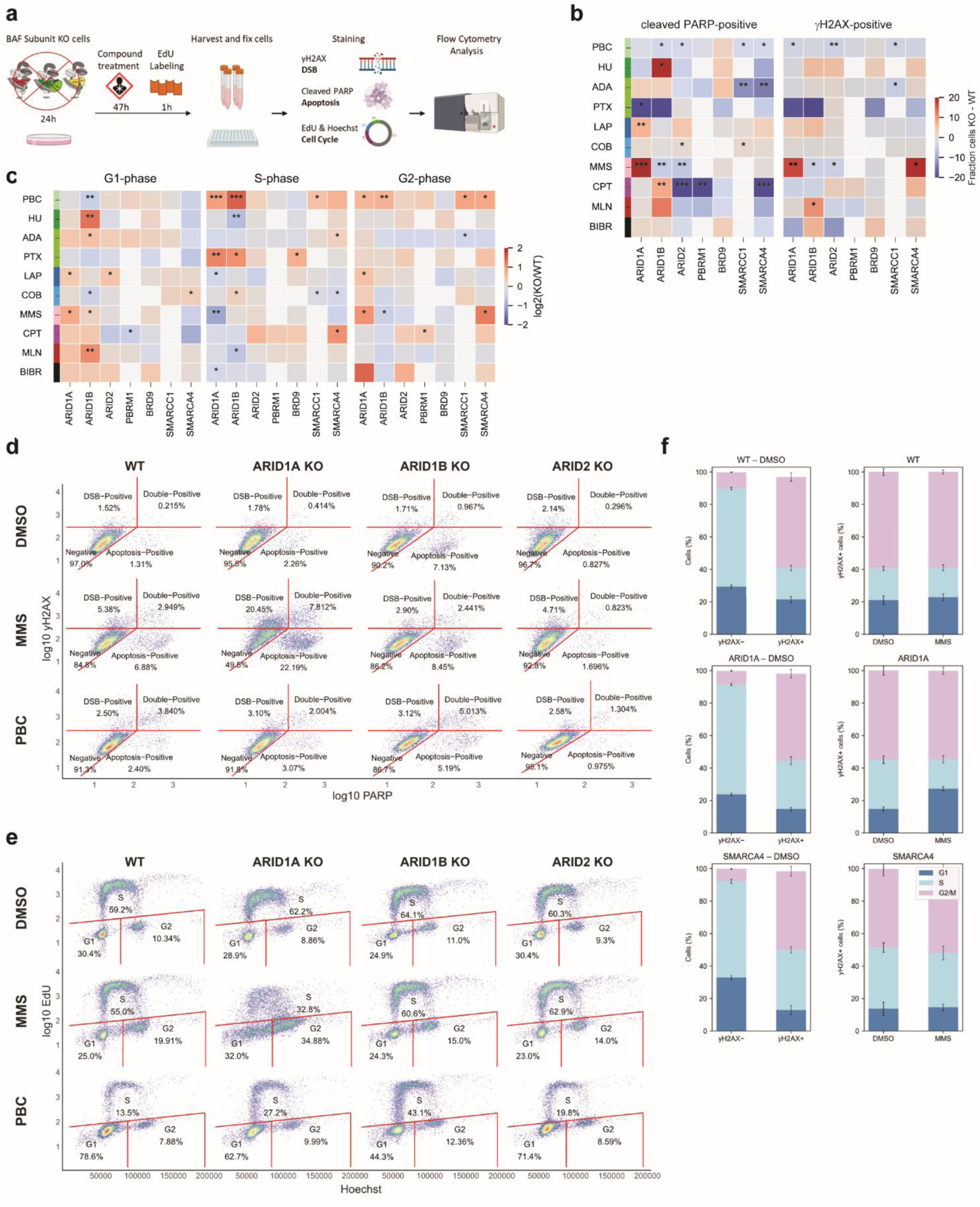
Cell cycle, apoptosis, and DSB alterations upon BAF KO and treatment. **a.** Schematic showing the flow cytometry screen setup. **b.** Heatmap showing difference in fraction of cleaved PARP (left) and γH2AX-positive cells (right) between BAF KO cells and WT subtracted each by the fraction of positive cells after treatment with DMSO. Measurements obtained from flow cytometry. Significance determined using t-test, p-values corrected for multiple testing, * p-value < 0.05, ** p-value < 0.01, *** p-value < 0.001, n≥3. PBC=Palbociclib, HU=Hydroxy Urea, ADA=Adavosertib, PTX=Paclitaxel, LAP=Lapatinib, COB=Cobimentinib, MMS=Methyl Methanesulfonate, CPT=Camptothecin, MLN=MLN4924, BIBR=BIBR1532. **c.** Heatmaps showing fraction of cells in G1 (left), S (middle), G2 (right) cell cycle phase in BAF KO cells upon treatment normalized to WT treated and each normalized to treatment with DMSO. Measurements obtained from flow cytometry quantifying EdU incorporation and Hoechst stain for DNA content. Statistics and abbreviations as in **b**. **d.** Dot plots obtained from flow cytometry stratifying cells by stains for cleaved PARP and γH2AX. Abbreviations as in **b. e.** Dot plots obtained from flow cytometry stratifying cells by Hoechst and EdU stain. Abbreviations as in **b. f.** Bar plots showing the fractions of cells in G1, S, and G2 phase for WT (top), ARID1A KO (middle), SMARCA4 KO (bottom) cells. Cells are split into γH2AX negative or positive cells upon treatment with DMSO (left). Bar plots to the right show γH2AX positive cells by cell cycle phase when treated with DMSO or MMS. Error bars indicate STE, n≥3. 4a) Created in BioRender. Bonn, L. (2026) https://BioRender.com/w78dkpf

As suspected from the cell growth measurements, we observed little to no significant signals in flow cytometry 48h after treating BAF KO cells with an EGFR or MEK inhibitor (Lapatinib and Cobimetinib, respectively), further supporting that sensitivities only manifest at later time points (Fig 4b-c). Similarly, treatment of BAF KO cells with MLN or BIBR1532 did not result in significant alterations in cell cycle phases, γH2AX or cleaved PARP levels, largely consistent with Incucyte measurements (Fig 4b-c). However, we observed significant reduction in apoptotic levels and less γH2AX-positive cells for various BAF KO cells upon treatment with the G1/S and G2/M checkpoint inhibitors Palbociclib, Adavosertib, and Paclitaxel already at the 48h time point (Fig. 4d), while in Incucyte measurements alterations in confluency mostly became significant at the 72h time point (Fig. 3d). Signals were especially strong in response to treatment with Palbociclib, for which the cBAF-specific ARID1A and ARID1B KO cells also displayed a significantly higher proportion of cells in G2 and S and less in G1 phase (Fig. 4e). This suggests that increases in cell growth are driven by both reduced apoptosis induction and faster cycling overcoming G1/S checkpoint inhibition by Palbociclib in comparison to WT. This effect seems to be specific for cBAF perturbation (Fig. 4d-e). Treatment with Paclitaxel resulted also in a higher proportion of cells in S phase for BRD9 KO cells, likely driving observed increases in cell growth in Incucyte measurements compared to WT (Fig. 3d, 4c, S4a). Conversely, Adavosertib treatment resulted in significantly less apoptotic cells in a SMARCC1 and SMARCA4 KO background while only SMARCA4 KO cells displayed increased cell growth (Fig. 3d, 4b-c, S4a). Hydroxy Urea (HU) treatment had caused significant increases in cell growth for SMARCA4, ARID1A and ARID2 KO cells at later time points but not for ARID1B (Fig. 3d, S4a). In flow cytometric analysis, we observed instead specifically for ARID1B KO cells higher levels of apoptosis and less cells in S but more in G1 phase (Fig. 4b-c). ARID1B KO cells might be sensitive to slowed replication as a consequence from HU treatment likely resulting from its accelerated growth behavior in untreated conditions in comparison to other BAF KO cell lines (Fig. 1b). In summary, while we confirm overall resistance to cell cycle checkpoint inhibition in BAF KO backgrounds, we discovered differences in cell cycle alterations and apoptosis induction depending on which checkpoint is blocked and which BAF subunit is perturbed.

DNA damaging agents showed significant effects on cell growth after 48h, in line with strong effects observed on cell cycle phases as well as apoptosis and γH2AX levels in flow cytometry measurements. Furthermore, these effects appear to be specific to individual BAF subunit perturbations. Treatment of cells with the DNA alkylating agents Methyl-Methanesulfonate (MMS) and IlludinS resulted in no significant alterations in cell growth during Incucyte measurements. However, we observed a non-significant yet specific sensitivity of ARID1A KO cells across time points when treated with either compound in contrast to slight resistances observed for ARID1B, ARID2, and SMARCA4 KO cells (Fig. 3d, S4a). In line with these observations, we observed strong apoptosis and γH2AX induction in ARID1A KO cells compared to reduced levels observed for ARID1B and ARID2 KO cells (Fig. 4b-d). Compared to SMARCA4 KO and WT cells, we observe that upon MMS treatment, ARID1A KO cells accumulate more γH2AX in G1 (Fig. 4f). Similarly, ARID1A but not ARID1B or ARID2 KO cells displayed fewer cells in S and more cells in G1 (also significant for ARID1B) and G2 phase (Fig. 4c,e). This suggests that ARID1A loss makes HAP1 cells particularly vulnerable to DNA alkylating agents, which cause DNA damage that, if left unrepaired, leads to replication fork stalling, collapse and DSBs during S phase. Unfortunately, we were unable to confirm with our confluency screen ncBAF subunit loss-specific effects upon treatment with IlludinS that were observed in the Olivieri dataset (Fig. 2b, 4b-c). Likewise, our cell growth and flow cytometric readouts did not confirm observed sensitivities of ncBAF perturbed cells when treated with the Top1 inhibitor Camptothecin, suggesting that these effects might be specific to the RPE-1 cell background (Fig. 3d, 4b-c). In summary, cell growth, cell cycle, cell fate and DNA damage phenotype measurements identified multiple unexpected sensitivities and resistances that are specific to individual BAF subunit or subtype loss upon treatment with growth signaling, cell cycle, or genome stability perturbants.

### Differential gene expression analysis generates mechanistic hypotheses underlying vulnerabilities upon BAF perturbation and treatment

To generate hypotheses about molecular mechanisms that might drive observed vulnerabilities via changes in gene expression programs, we performed total transcriptome analysis via RNA-seq on SMARCA4, ARID1A/B, ARID2, and BRD9 KO cells as well as WT cells when treated for 48h with Camptothecin, Adavosertib, MMS, Palbociclib, MLN, or HU (Table S6). Generally, treatment of BAF KO cells with HU, MLN, and MMS showed much more mild effects on gene expression compared to treatment with Palbociclib, Camptothecin, and Adavosertib (Fig. 5a). Clustering of genes that are differentially expressed between treated BAF KO and WT (each compared to DMSO), revealed groups of genes with interesting functional enrichments (Fig. 5a, Table S6). Cluster 1 is highly enriched for upregulated genes, especially for ARID1A, ARID1B, and SMARCA4 KO cells treated with Palbociclib, that function in sister chromatid segregation, spindle assembly, and mitotic cell phase transition (Fig. 5a), which is in line with observations from flow cytometry and partially from Incucyte measurements that showed increased cell growth, less apoptosis, and a shift of cells towards S and G2 phase. Upon closer inspection we find upregulation of most MCM complex members, cyclins, checkpoint proteins (BUB1, BUB1B, Chk2) as well as centromeric and kinetochore related proteins (Fig. S5a). Cluster 2 consists of downregulated genes primarily from ARID1B and BRD9 KO cells when treated with Adavosertib, Camptothecin, MMS, and Palbociclib that function in amino acid import and tRNA aminoacetylation as well as in responses to ER stress and starvation (Fig. 5a). Genes regulating cell growth and size as well as components of the Wnt and Notch signaling pathway are the most downregulated in ARID1B KO cells upon treatment with Palbociclib (cluster 3) (Fig. 5a). Of interest is also cluster 5 showing an enrichment for genes functioning in DSB repair via HR, DNA replication, checkpoint signaling and negative regulation of cell cycle phase transitions (Fig. S5b). These genes are primarily downregulated in SMARCA4 KO cells treated with Camptothecin potentially driving its reduced apoptosis levels observed in flow cytometry (Fig. 4b, S5b). Upon closer inspection we find that ARID1A/B and SMARCA4 KO cells treated with Palbociclib show an upregulation of members of base excision repair (APEX1, NEIL2, MPG, NTHL1, PNKP), nucleotide excision repair (ERCC1/2, RAD23A), homologous recombination (FANCD2, ABRACAS1, CHEK2, SPIDR, RMI2, RAD51, BRCA1) and Fanconi Anemia pathway compared to treated WT and DMSO controls (Fig. S5b). Contrary to that, genes functioning in cell cycle regulation such as CDK1, PLK1, AURKB, WEE1, CHEK2, and ATR as well as members of the anaphase promoting complex and its regulators are downregulated in ARID1A KO cells when treated with MMS compared to treated WT and DMSO controls (Fig. S5a), potentially driving observed sensitivities in flow cytometric analysis.

**Figure 5:**
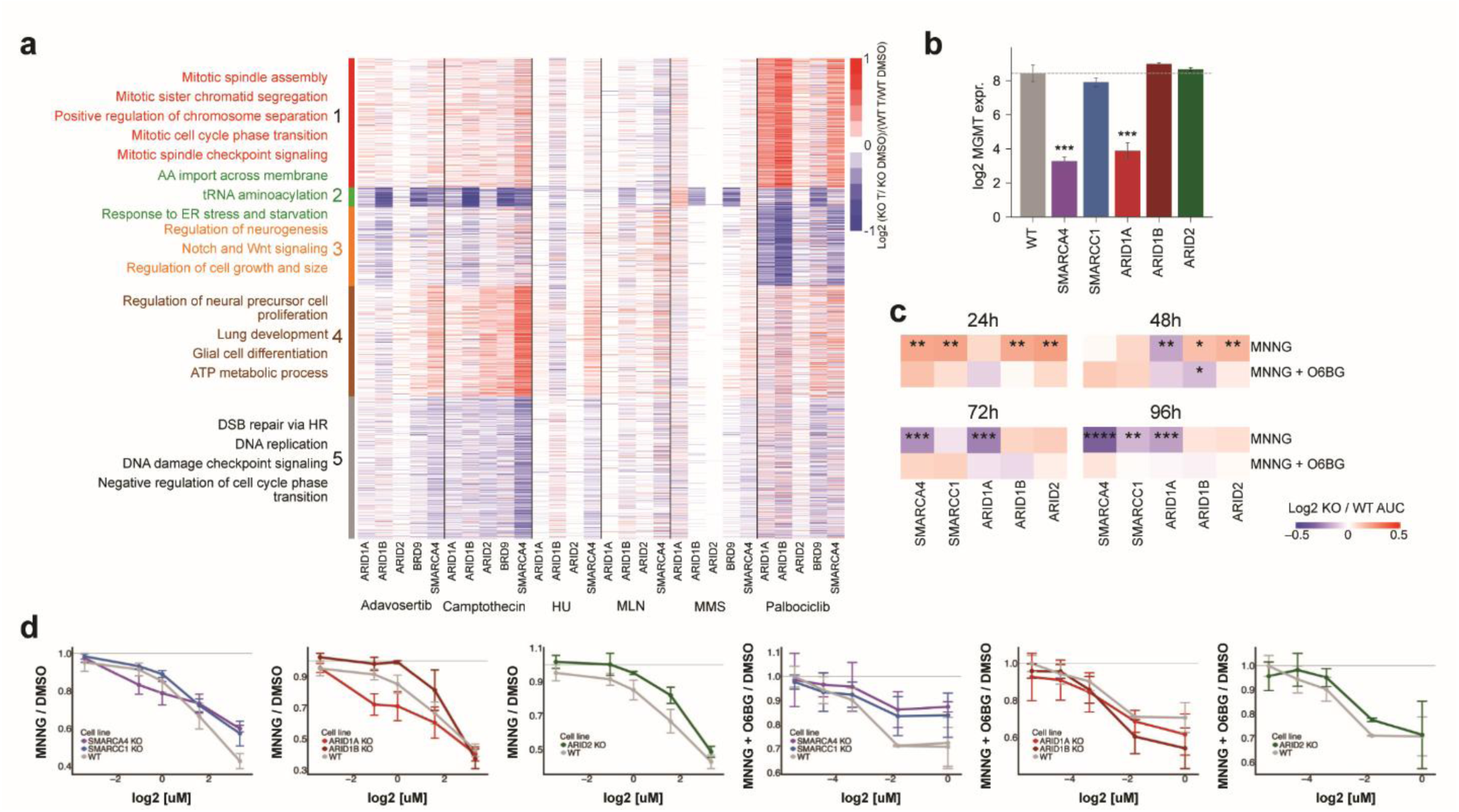
Transcriptome analysis reveals BAF KO gene expression changes driving compound sensitivities and MGMT-driven MNNG vulnerability. **a.** Heatmap showing clustered differentially expressed genes from total transcriptome analysis in BAF KO vs WT cells 48 h after treatment with indicated compounds normalized to DMSO-treated controls. Clusters were analyzed for enriched GO terms. Representative enriched GO terms are indicated on the left. HU=Hydroxy Urea, MMS=Methyl Methanesulfonate, MLN=MLN4924. n=3. **b.** Bar plot showing MGMT expression levels for selected BAF KO cells and WT. **c.** Heatmaps showing area under the dose response curves (AUC) from confluency measurements for BAF KO over WT upon MNNG ± O6BG treatment and normalized to DMSO controls. Significance determined using t-test, p-values corrected for multiple testing, * p-value < 0.1, ** p-value < 0.05, *** p-value < 0.01, **** p-value < 0.001, n=3. MNNG = Methylnitronitrosoguanidine, O6BG = O6-Benzylguanine. **d.** Dose response curves of confluency measured 48 h after treatment for various BAF KO cells and WT upon MNNG ± O6BG treatment normalized to treatment with DMSO. Error bars indicate STD, n=3.

Since we discovered striking differential expression of MGMT across several BAF KO cells in untreated conditions in comparison to WT (Fig. 1l, 5b) and a further downregulation of MGMT expression levels in ARID1A KO cells upon treatment with Adavosertib, CPT, or HU while MGMT was upregulated in SMARCA4 KO cells upon treatment with Camptothecin (in comparison to WT treated and normalized to DMSO) (Fig. S5b), we set out to investigate to which extent MGMT expression levels are predictive of sensitivities or resistances of BAF KO cells upon treatment with MNNG, a DNA alkylating agent whose damage is reversed by MGMT^30^. Since any given MGMT molecule is inactivated upon the reversal reaction, we expect that MGMT transcript levels should correlate well with cell survival upon treatment with MNNG. Using our live-cell imaging setup we indeed observed that SMARCA4 and ARID1A KO cells with reduced MGMT expression levels were particularly sensitive to MNNG treatment (Fig. 5c-d). This sensitivity was specific to MGMT activity because treatment with the MGMT inhibitor O6BG abrogated these sensitivities (Fig. 5c-d). This highlights how detected changes in gene expression levels might point to mechanisms driving observed differential phenotypes upon BAF perturbation.

## Discussion

The frequent mutations of genes encoding subunits of the BAF chromatin remodeling complexes across diverse cancers have raised interest in understanding how loss of individual subunits affects gene regulation, genome stability, and cellular fitness. Several studies have made interesting findings for several subunits across different conditions. However, a comprehensive understanding has been complicated by the strong context dependence of BAF complex activity and by the combinatorial assembly of its subunits into distinct complexes. To systematically disentangle subunit- and subtype-specific effects, we employed an isogenic panel of HAP1 cell lines comprising knockouts of key subtype-defining and common BAF complex subunits. This approach enabled direct comparison of cellular phenotypes and drug responses across the same genetic background.

Our results reveal that disruption of individual BAF subunits produces distinct and sometimes opposing effects on cell proliferation, genome stability, and drug sensitivity. Loss of the two most frequently mutated subunits in cancer, ARID1A and SMARCA4, did not accelerate proliferation but resulted in elevated γH2AX levels and a higher maximum capacity. This indicates increased DNA damage burden and reduced density-dependent growth inhibition – characteristics often observed in cancer cells. Notably, γH2AX accumulation in SMARCA4-deficient cells was particularly pronounced during S and G2/M phase, in agreement with previous reports demonstrating a role of SMARCA4 in homologous recombination-mediated DSB repair during these stages of cell cycle^18,31^. The higher maximum capacity may reflect altered contact inhibition, density-dependent signaling, metabolic tolerance or cell-cell interaction. Consistently, transcriptome measurements of SMARCA4 KO cells revealed upregulation of adhesion-related genes and multiple growth signaling genes.

In contrast, knockout of ARID1B had a markedly different phenotype characterized by increased proliferation without a change in maximum capacity. This divergence between ARID1A and ARID1B supports the emerging view that these paralogous subunits are not functionally redundant but instead mediate distinct regulatory functions within cBAF complexes. Consistent with our observation, ARID1B loss has been shown to promote leukemogenesis in mouse models driven by the MLL-AF9 fusion^32^, a context mechanistically related to the BCR-ABL fusion present in HAP1 cells^33^. Conversely to ARID1B KO cells, knockout of the core scaffold subunit SMARCC1 caused the most severe growth impairment, with reduction in both growth rate and maximum capacity. The associated transcriptional upregulation of cell adhesion and developmental genes suggests that SMARCC1 loss may promote differentiation-like states or reinforce contact inhibition pathways, thereby limiting proliferative potential. Together, these findings highlight the functional heterogeneity of BAF subunits and demonstrate that disruption of different components can differently influence proliferation and genome stability.

Consistent with these diverse cellular phenotypes, our compound screens revealed both shared and subunit-specific responses to geno- and cytotoxic agents. Unexpectedly, many BAF knockout lines exhibited reduced sensitivity to several DNA-damaging drugs and cell cycle checkpoint inhibitors compared to wildtype cells. This trend contrasts with the prevailing model that loss of BAF subunits impair DNA repair and sensitizes cells to genotoxic stress. However, a few examples of drug resistance have been reported previously, including resistance of ARID1A- or SMARCB1-deficient cells to the anthracycline Doxorubicin^34,35^ and of SMARCB1-deficient cells to Paclitaxel^35^. This is in line with our observed resistance of ARID1A knockout cells to Doxorubicin and Paclitaxel. Our findings extend these observations by showing that multiple BAF perturbations can confer resistance to a broader range of genotoxic agents.

The pronounced resistance of SMARCA4 KO cells to Top1 inhibitors may arise from impaired chromatin engagement of Topoisomerase I, as SMARCA4 has been shown to facilitate its recruitment^36^. Despite an unchanged fraction of cells in S-phase, SMARCA4 knockout cells exhibited elevated DNA damage specifically during DNA replication, indicative of replication stress. Slower or stalled replication forks could reduce collisions between replication machinery and Topoisomerase I cleavage complexes stabilized by the inhibitors, thereby limiting cytotoxic double-strand break formations and conferring drug resistance. In addition, previous studies in SMARCA4-deficient cancer cells have reported dysregulation of apoptotic signaling pathways^37^, which may further contribute to reduced sensitivity to cyto- or genotoxic treatments.

The strongest resistance in our growth screen was observed for the PARP inhibitor Niraparib, which inhibits PARP catalytic activity and traps PARP-DNA complexes^38^, thereby leading to cytotoxic DNA lesions. Resistance was particularly pronounced in knockouts of the subtype-defining ARID subunits as well as the common subunits SMARCA4 and SMARCC1. This contrasts with reported sensitivity of ARID1A-deficient cells to PARP inhibition due to impaired DNA damage checkpoint signaling^20,39^. Notably, analysis of the PRISM dataset likewise revealed a general resistance to Niraparib across various BAF deficiencies and cell types, arguing that this phenotype may represent a broader feature of BAF deficiencies. One possible explanation is that altered chromatin accessibility or replication dynamics in BAF-deficient cells reduce the formation or cytotoxic consequences of PARP-trapping lesions. Alternatively, compensatory rewiring of DNA repair pathways or increased tolerance to DNA damage may contribute to this phenotype. Consistent with this interpretation, BAF knockout cells in our screen also displayed reduced sensitivity to several other DNA damage-inducing compounds.

Resistance to the CDK4/6 inhibitor Palbociclib was observed in common and particularly cBAF-specific knockouts. Loss of ARID1A and ARID1B drives CCNE1 upregulation after Palbociclib treatment, sustaining Cyclin E/CDK2 activity and bypassing CDK4/6 inhibition to confer resistance. In contrast, SMARCA4-deficient non-small cell lung cancer and hypercalcemic small cell carcinoma of the ovary exhibit increased Palbociclib sensitivity due to restricted chromatin accessibility at the *CCND1* locus and reduced Cyclin D1 expression^40^. In our system, knockout of the ncBAF-specific subunit BRD9 and SMARCD1 – the only SMARCD subunit in ncBAF – resulted in the lowest CCND1 expression across all tested HAP1 cells and upregulation upon Palbociclib treatment, resulting into a trend towards sensitivity. Notably, ARID2 and SMARCA4 knockouts, initially less sensitive, displayed a pronounced retraction after 48 to 72 hours, suggesting that the responses to cell cycle-targeting drugs may emerge more gradually than those of genotoxic agents and require longer observation windows.

In contrast to the predominantly resistant phenotypes in BAF knockout cells observed with cyto- and genotoxic agents, inhibitors targeting growth signaling _-_particularly EGFR and MEK _-_showed an overall trend towards increased sensitivity in BAF-deficient cells. Analysis of the PRISM dataset supported this observation and suggested that the effect is most pronounced in perturbations affecting PBAF subunits: Cells with low PBRM1 expression showed increased sensitivity to EGFR inhibitors, while reduced ARID2 expression was associated with sensitivity to MEK inhibition. Similar trends were observed for reduced expression of other BAF subunits, particularly ARID1A and ARID1B. Notably, HAP1 ARID1B knockout cells exhibited increased resistance to EGFR inhibition at later timepoints, consistent with their overall proliferative advantage and highlighting the functional heterogeneity of BAF subunits.

The strongest sensitivity was observed for the MEK inhibitor Cobimetinib in SMARCC1 knockout cells. Similar to SMARCA4-deficient cells, these cells have elevated expression levels of AP-1 target genes, including FOS, JUN, EGR1, suggesting increased reliance on AP1-driven transcriptional programs downstream of ERK signaling to sustain proliferation. Such transcriptional rewiring is consistent with the role of BAF complexes in regulating enhancer accessibility. Perturbation of BAF complex function may therefore shift proliferative control toward EGFR-MAPK-dependent transcriptional circuits, creating signaling dependencies that can be therapeutically exploited. Indeed, activation of EGFR-MAPK signaling has been reported in several BAF-deficient contexts^16,41,42^. Interestingly, these effects became most evident at later timepoints in our assay, coinciding with the transition out of exponential growth. This observation raises the possibility that BAF-deficient cells increasingly rely on mitogenic signaling to overcome density-dependent growth arrest.

Our data also revealed subunit-specific responses to DNA repair-targeting agents. ARID1A knockout cells were uniquely sensitive to the DNA alkylating agent MMS and showed induction of DNA damage signaling and apoptosis. This phenotype is consistent with the reported role of ARID1A in base excision repair (BER), where its loss delays recruitment of BER repair proteins and leads to accumulation of base lesions and abasic site^43^. Notably, this defect was not observed upon loss of its paralog ARID1B or the PBAF-specific homolog ARID2. We observed several of these distinct effects of different ARID knockouts also in other contexts, such as increased proliferation, selective resistance to Palbociclib and sensitivity to Hydroxy Urea in ARID1B knockouts. These findings reinforce the concept that individual ARID subunits exert nonredundant functions within distinct BAF complexes. Indeed, we reported previously that ARID1A loss causes widespread loss of chromatin accessibility, whereas ARID1B loss results in increased accessibility at many genomic regions^8^. Moreover, while overall differentially expressed genes are similar between ARID1A and ARID1B loss, ARID1B knockout cells showed stronger downregulation of growth signaling factors and more differentially expressed DNA damage response genes compared to ARD1A and ARID2 knockouts.

We were able to confirm previously observed sensitivity of PBAF-deficient cells to the NEDD8-activating enzyme (NAE) inhibitor MLN4924, and increased sensitivity of ARID2- and PBRM1-deficient cells to EGFR/MEK inhibitors. At the same time, certain effects appear to be context-dependent. For example, ncBAF-specific sensitivities to illudinS observed in RPE-1 CRISPR screens^28^ were not observed in our HAP1 model and HAP1 showed a hypersensitivity to Topoisomerase I inhibitors compared to RPE-1 cells. Apart from the influence of cellular context, additional differences between studies included the use of stable knockouts versus acute CRISPR perturbations and variations in treatment duration or dosing regimes, which may also influence the observed phenotypes.

Our study also has several limitations. All experiments were performed in a single cellular background, raising the possibility of cell type- or clone-specific effects. Moreover, stable knockout cells may have adapted to the loss of BAF components through compensatory transcriptional or signaling changes. Future studies across additional cellular contexts and using complementary perturbation strategies will therefore be important to further investigate the identified vulnerabilities. Longer treatment windows may also be necessary to fully capture dependencies on growth signaling pathways that emerge outside of exponential proliferation.

Overall, our systematic analyses uncovered vulnerabilities of selected BAF-deficient cells to MEK and EGFR pathway inhibition, as well as resistance toward checkpoint inhibitors and certain DNA-damaging agents. We identified clear BAF subunit-dependent effects, underlying their specific contributions to cell cycle and genome maintenance regulation.

Taken together, our findings support a model in which disruption of BAF chromatin remodeling complexes rewires transcriptional and signaling networks that control proliferation, genome maintenance, and drug response. Loss of key BAF subunits leads to context-dependent shifts in regulatory programs governing cell cycle progression, DNA repair, and growth signaling. Distinct BAF subunits contribute specialized regulatory functions: the catalytic subunit SMARCA4 maintains chromatin states that support efficient DNA repair and replication, subtype-defining ARID proteins fulfil non-redundant functions in chromatin regulation and genome maintenance, and scaffold subunits such as SMARCC1 stabilize complex integrity and proliferative programs. Disruption of these functions produces heterogeneous cellular outcomes but converges on three interconnected regulatory layers: genome maintenance, cell cycle control, and mitogenic signaling. These alterations can manifest as replication stress and basal DNA damage, rewiring of cell cycle transcriptional programs that e.g. bypass canonical CDK4/6 control, and increased reliance on EGFR-MAPK-dependent proliferative signaling, thereby creating targetable vulnerabilities. This combination may help explain the coexistence of chemoresistance and sensitivities in distinct BAF-mutant cells and suggests that specific BAF chromatin remodeler defects can rewire cellular fitness to create both therapeutic liabilities and opportunities.

## Acknowledgements

We thank all members of the Schick and Luck lab for helpful discussions throughout the study. We thank Magdalena Schachtl-Rieß for help with cloning and Leon Maximillian Persch for help with DepMap data analysis. We thank the Christmann lab for protocols and advice on MNNG treatments. We thank the IMB Microscopy and Histology CF, especially Sandra Ritz and Rossana Piccinno, for assistance with high-throughput confocal imaging on the Opera Phenix (DFG project #316215830) and with live-cell imaging on the Incucyte. We thank the IMB Flow Cytometry Core Facility, Stefanie Möckel and Stephanie Nick, for their assistance with and for usage of the BD LSRFortessa SORP (DFG project #210253511), BD FACSAria III (DFG# 210144599) and Invitrogen Bigfoot (DFG #511658729). We thank the IMB Genomics Core Facility for assistance with the NGS experiments and the use of the NextSeq2000. We thank the Media Lab facilities from IMB. The research performed in this study was funded by the Deutsche Forschungsgemeinschaft (DFG, German Research Foundation) – Project-ID 393547839 – SFB 1361 awarded to KL and SS, Project-ID 449991970 awarded to KL.

## Author contributions

K.L. and S.S. designed and coordinated the study, K.S., C.N., M.Wo., L.B., M.We., and C.S. performed experiments and analyses, C.B., E.A., F.R. and E.S. performed experimental data analysis, C.B., E.A., and E.S. performed computational work, K.S., C.B., C.N., E.A., K.L. and S.S. wrote the manuscript and K.L. and S.S. provided the funding.

## Data and code availability

The GrowPipe package for the Incucyte data analysis is available at https://github.com/KatjaLuckLab/GrowPipe_public. Code for all other data analysis is available at https://github.com/KatjaLuckLab/BAF_compound_screen_paper.

## Methods

### Cell lines and cell culture

#### Cell culture

In this study, we used a selection of HAP1 WT and BAF mutant HAP1 cell lines (single–gene knockouts for individual BAF subunits) described previously (Schick et al., Nat Genet 2019). The cells were cultured in IMDM (Thermo Fisher Scientific, Cat.# 12440061) supplemented with 10% Value Fetal bovine serum (VFBS) (Thermo Fisher Scientific, Cat.# A5256701) and 1% penicillin-streptomycin (5000 U/ml, Thermo Fisher Scientific, Cat.# 15070063). The cells were incubated under standard conditions at 37 °C, 5% CO₂ and 85% humidity. Subculturing was performed every 2–3 days when the cells reached approximately 70–80% confluence.

#### Selection for diploid HAP1 KO cells

For this study, we used diploid HAP1 cell lines, as diploid cells more closely resemble the ploidy of most human somatic cells and exhibit improved stability in long–term culture compared to haploid HAP1 cells. Because HAP1 cells spontaneously diploidize over time, diploid lines were isolated by Fluorescence-Activated Cell Sorting (FACS) using the BD FACSAria III or Invitrogen Bigfoot cell sorter using a 100 µm nozzle and 20 psi setup in purity sort precision mode. In brief, cells were gated by forward/side scatter (FSC-A vs SSC-A) for size/granularity, singlets were selected, dead cells excluded by DAPI staining, and live single diploid (DAPI-negative) cells collected and cultured for downstream experiments (Fig S1a). The ploidy status was verified by propidium iodide staining followed by cell-cycle analysis (Fig. S1b, c).

### Compound treatments

### Live cell growth screen

To determine the growth rate in dependency of genotype and treatment, the confluency of cells were monitored over several days by taking images of the cells with the Incucyte® SX5 Live-Cell Imaging and Analysis System (Sartorius, Göttingen, Germany), a microscope integrated into an incubator that allows the cells to grow undisturbed while being imaged.

### Poly-D-lysine (PDL) coating of plates

50 μl PDL (50 μg/ml in DPBS, Merck, Cat.# A-003-E) per well was incubated for 10 min at room temperature (RT) on an orbital shaker (30 rpm). PDL solution was removed and 100 μL DPBS (Thermo Fisher Scientific, Cat.# 14190144) added for plate storage at 4 °C (≤ 1 week). Before use, DPBS was removed and plates air-dried for one to three hours with the lid open in a laminar flow cabinet.

### Seeding of cells

The cells were split one day prior to seeding to reach 50-60% confluency in the tissue culture (TC) plate at seeding. On the day of seeding, the cells were washed with DPBS and detached with 0.25% trypsin/EDTA (Thermo Fisher Scientific, Cat.# 25200-056) for 5 min at 37°C. The trypsin reaction was stopped by adding 4x the volume of fresh culture medium. The cell suspension was transferred to a 15 ml tube and centrifuged at 300×g for 5 min. The resulting cell pellet was resuspended in fresh medium and counted twice with the Celldrop FL CellCounter. Based on the mean of these measurements, the cell suspension was diluted to 1*10^5^ cells/ml. This cell solution was counted four times with the Celldrop FL CellCounter, and then further diluted to an appropriate concentration. Unless otherwise stated, a concentration of 3*10^4^ cells/ml was created to seed 4500 cells in 150 µl of medium per well. The cells were pipetted onto Poly-D-lysine coated Corning® 96-well Clear Flat Bottom Polystyrene TC-treated Microplate (Corning, USA) and left for 30 min at RT to allow the cells to settle to the bottom of the well. Subsequently, the cells were placed in the Incucyte® SX5 and further incubated at 36.5°C, 5% CO2 and 85 % humidity for up to 7 days. Here, every 6h, 5 phase images per well were taken with a 10x magnification, using the Adherent Cell-by-Cell scanning mode (image resolution = 1.24 µm/pixel; image size = 1408 × 1040 pixels; field of view = 1.75 × 1.29 mm).

### Treatment of cells

If compounds were used, the cells were treated 24h after seeding. For this, the compounds were diluted from their stock solutions to the final dilution (Table 1) in fresh cell culture medium. DMSO concentrations were adjusted to be the same for all treatments with the same compound. The compound containing medium was prewarmed for 30min. The cell plates were removed from the Incucyte 20min before the 24h timepoint. 10min before the 24h timepoint, the old medium was removed from the plate and exchanged against fresh medium containing compounds. 5min before the 24h timepoint, the plate was placed back in the Incucyte® SX5 and then imaged for the first time with the compound 24h after seeding. After that, the plate was imaged every 6h for up to 5 days.

**Table 1.**
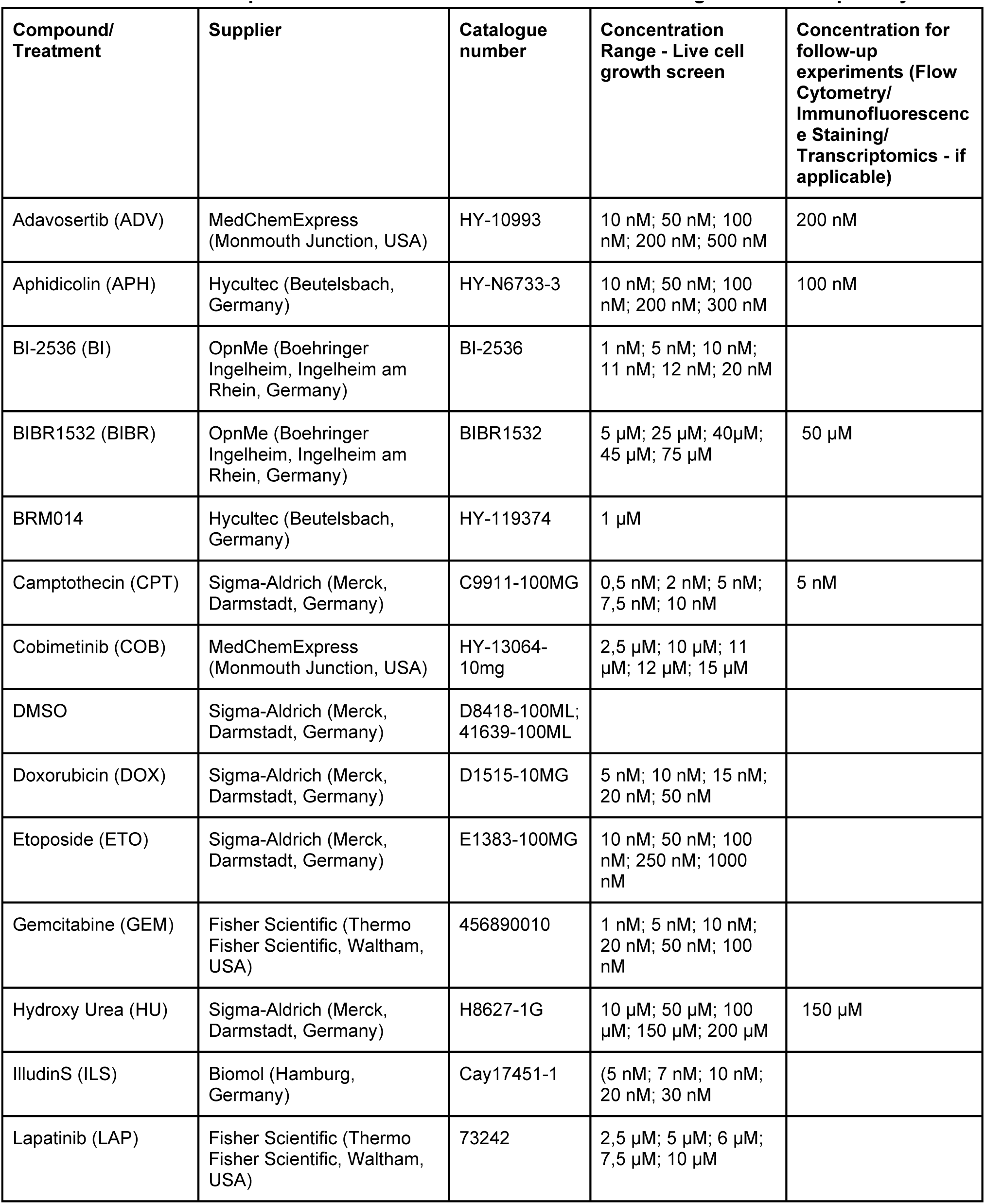
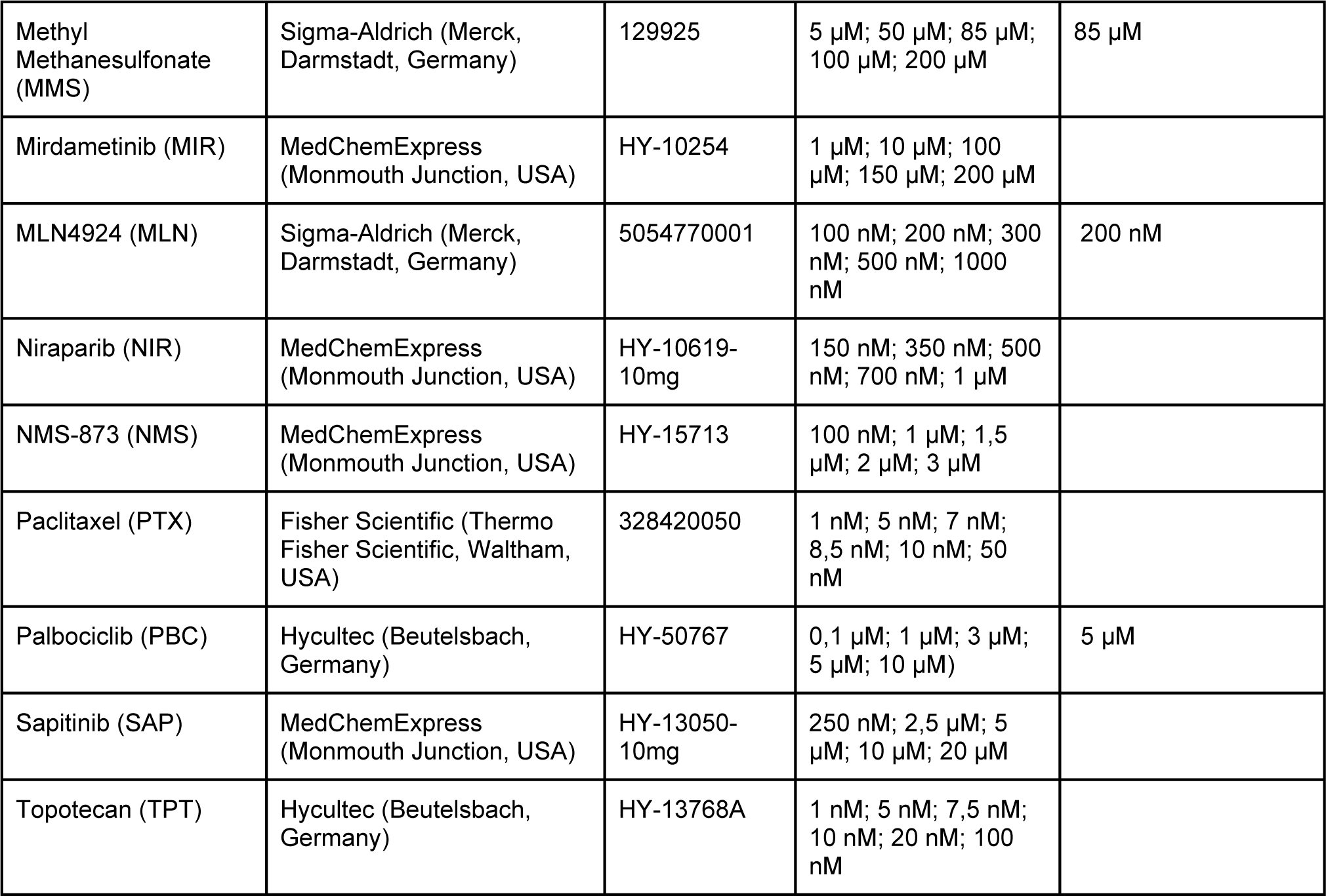
Overview of Compounds and Concentrations used for Screening and Follow-up assays.

### Analysis of Confluency

The confluency of the images taken by IncuCyte was determined with the IncuCyte software (Essen BioScience Inc., software v2021A) using the Basic Analyzer. The segmentation adjustment was set to 1.1, the hole fill was set to 0, and the adjusted size of the pixels was -2. Objects with an area under 90 µm^2^ were filtered out. For each time point, 5 images (fields of view) were taken per well, and the confluency per well corresponds to the mean confluency across these 5 images.

### Flow cytometry screen

Changes in cell cycle phase distribution as well as γH2AX and cleaved PARP levels upon treating cells with genotoxic compounds were measured using flow cytometry (FC).

### Seeding and treatment of cells

A suspension of the cells were prepared as described for the live cell growth screen and cell concentration measured using the Celldrop FL CellCounter. Each cell line was counted three times, and the average cell count was calculated. Following this, 3.6*10^5^ cells were seeded into 6 cm CytoOne® TC dishes (STARLAB International, Hamburg, Germany) in 3 ml of culture medium and incubated for 24 h under standard conditions. Cells were then treated by replacement of the medium to 4 ml of fresh medium with compound and further incubated. All compounds were first pre-diluted in DMSO and then diluted to the final concentration (Table 1) in culture medium, to ensure the same final DMSO concentration across all treatments. A separate DMSO control was included for each group of samples processed at the same time. Each treatment was then compared to its corresponding DMSO control.

### Click-iT^TM^ EdU Assay including γH2AX- and cleaved PARP staining

We used a Click-iT™ Plus EdU Alexa Fluor™ 488 Flow Cytometry Assay Kit (Invitrogen, Thermo Fisher Scientific, Waltham, USA, Cat.# C10633), with additional staining for γH2AX (Merck, Darmstadt, Germany) and cleaved PARP (Cell Signaling Technology, Cambridge, UK) to evaluate the cell cycle stage, DNA damage, and apoptosis in cells treated with genotoxic agents. The assays were done according to the manufacturer’s protocol with some adjustments, as the measurements were performed as a high-throughput screen in a 96-well format.

### Labeling cells with EdU

Cells were labelled with 5-Ethynyl-2‘-deoxyuridine (EdU) 1 h prior to harvesting as described in the manufacturers protocol by adding 4 µl of the 10 mM stock solution directly into the cell culture medium on the plates (final concentration: 10 µM). The plates were carefully shaken to evenly distribute the EdU and incubated under standard conditions for one hour.

### Harvesting and fixation of cells

Cells were harvested 48h post treatment. To capture dead cells in the medium, medium was collected and combined with the harvested cells after trypsinization. The tubes were centrifuged at 300×g for 5 min. The pellet was then resuspended in 250 µl of 1% BSA in PBS solution and transferred to a 96-well microplate with V-shaped bottoms (Starlab, Hamburg, Germany, Cat.# S1833-9600). The cells were washed once more with 1% BSA in PBS before being resuspended in 100 µl of Click-iT fixative, which consists of 4% paraformaldehyde, and incubated at room temperature for 15min. Afterwards, the cells were washed twice with 1% BSA in PBS before being permeabilized in 100 µl of Click-iT wash and permeabilization solution, followed by another 15 min incubation. Finally, the cells were resuspended in 250 µl of 1% BSA in PBS.

### Counting Cells with the Incucyte® Live-Cell Imaging and Analysis System

To better handle the large sample size, a method was developed that makes it possible to effectively and quickly determine the cell concentration of the individual samples using the Incucyte® Live-Cell Imaging and Analysis System. A small volume of cell suspension (1 µl) was transferred to a Corning® 96-well Clear Flat Bottom Polystyrene TC-treated microplate (Corning, Corning, USA, Cat.# 3598) containing 100 µl of PBS per well, in duplicates using a multichannel pipette. To ensure even distribution of the cells, the cell suspension was resuspended by pipetting and the plate was centrifuged at 1000×g for 5 min before being imaged using the Incucyte®. Nine images per well were taken and the average cell count per image was estimated using the Incucyte® analysis system. The average cell count per image was then multiplied by nine to obtain the total number of cells within the captured area. Since the well area was 6,089 times larger than the nine imaged areas combined, the calculated cell count was multiplied by 6,089 to estimate the total number of cells per well. To determine the cell concentration in suspension, this value was then multiplied by 1,000, yielding the number of cells per milliliter. Finally, the required volume of cell suspension for labeling 300,000 cells was calculated.

### Detect EdU – Click reaction

A total of 300,000 cells per sample were transferred to a new 96-well microplate with V-shaped bottoms. The cells were then centrifuged at 1000×g for 10 min, the supernatant was carefully removed, and the cells were resuspended in 50 µl of Click-iT wash and permeabilization solution. The EdU buffer additive was freshly prepared by diluting the 10× stock solution to 1× in deionized water. Subsequently, the Click-iT Plus reaction cocktail was prepared according to the manufacturer’s instructions.

As this high-throughput screen was performed in a 96-well format, the required volumes were downscaled. For staining 300,000 cells, 150 µl of the cocktail was added to each sample. The components used in the screen and their adjusted volumes are listed in Table 2. The cells were incubated on a shaker (Polymax 1040, Heidolph, Schwabach, Germany) at 30 rpm protected from light for 30 min and afterwards washed with 200 µl of Click-iT wash and permeabilization solution.

**Table 2.**
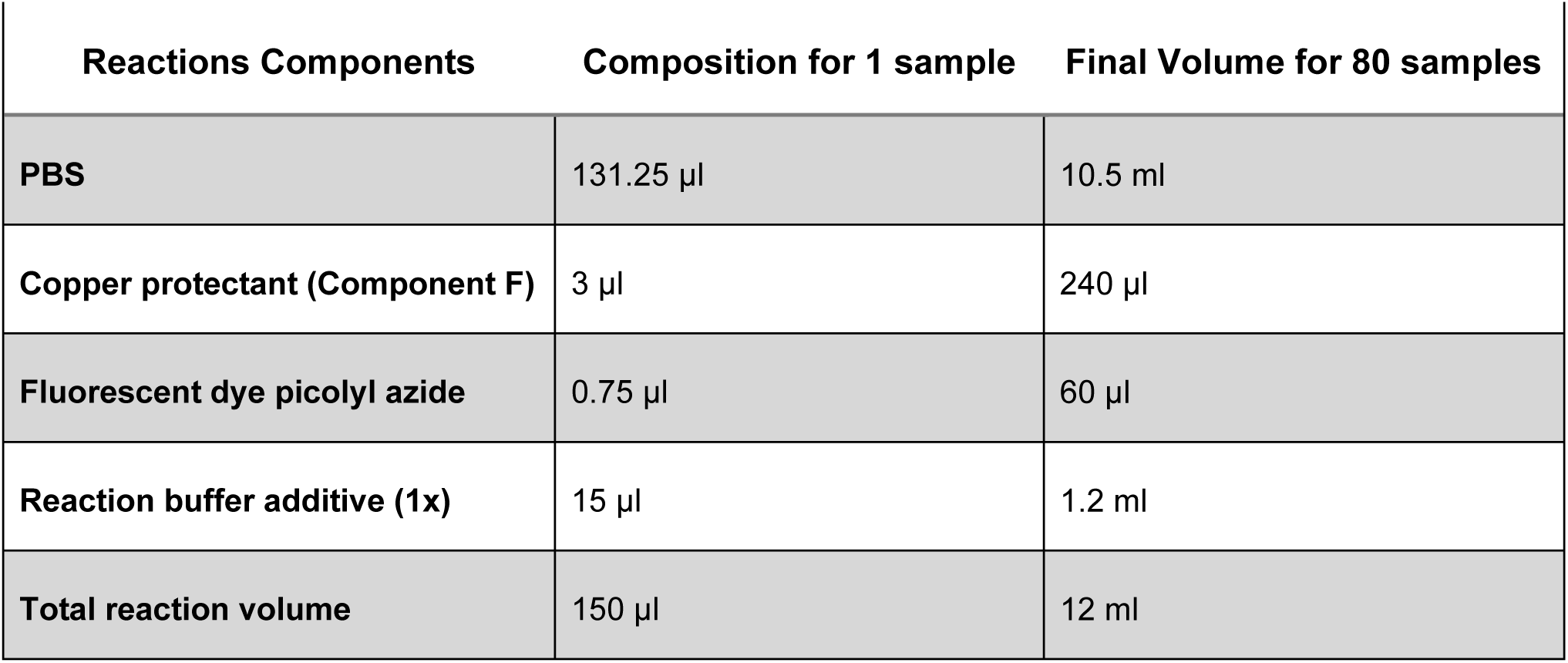
Reaction Cocktail Composition Used in the Flow Cytometry High-Throughput Screen.

| Reactions Components | Composition for 1 sample | Final Volume for 80 samples |
| --- | --- | --- |
| **PBS** | 131.25 μl | 10.5 ml |
| **Copper protectant (Component F)** | 3 μl | 240 μl |
| **Fluorescent dye picolyl azide** | 0.75 μl | 60 μl |
| **Reaction buffer additive (1x)** | 15 μl | 1.2 ml |
| **Total reaction volume** | 150 μl | 12 ml |

### Staining for DNA Damage and Apoptosis

The antibodies γH2AX and cleaved PARP (Table 3) were incubated together in 100 µl of 1% BSA in PBS on a shaker at 30 rpm, protected from light, for 1h. Following incubation, the cells were washed with 200 µl 1% BSA in PBS and resuspended in 200µl Click-iT wash and permeabilization solution. Finally, the DNA was stained with 0,002 mg/ml Hoechst 33342 (Life Technologies, Thermo Fisher Scientific, Waltham, USA, Cat.# H1399). All samples were analyzed using the High Throughput Sampler (HTS) of the BD LSRFortessa SORP (BD Biosciences, Franklin, USA) within two hours after staining.

**Table 3.** List of antibodies used for the Flow Cytometry screen.

| Antibody | Conjugated<br>Flourophore | Species<br>of Origin | Clone | Vendor | Cat. # | Lot. # | Dilution |
| --- | --- | --- | --- | --- | --- | --- | --- |
| **γH2AX**<br>**(pS139)**<br>**AF647** | Alexa 647 | Mouse<br>IgG1 | N1-431 | BD Biosciences<br>(Franklin, USA) | 560447 | 3247924 | 1:80 |
| **Cleaved**<br>**PARP**<br>**(Asp214)**<br>**PE** | PE | Rabbit<br>IgG | D64E10 | Cell Signaling<br>Technology<br>(Cambridge, UK) | 8978S | 2 | 1:250 |

### Measuring Cells with Flow Cytometry

Since multiple fluorophores were used to stain different antigens simultaneously, it was essential to ensure that fluorescence intensity peaks were sufficiently distinct to avoid bleed-through. To achieve this, the Fluorescence SpectraViewer (Thermo Fisher Scientific, Waltham, USA) was used to design a flow cytometry panel with lasers and their corresponding filters, as described in Table 4.

**Table 4.**
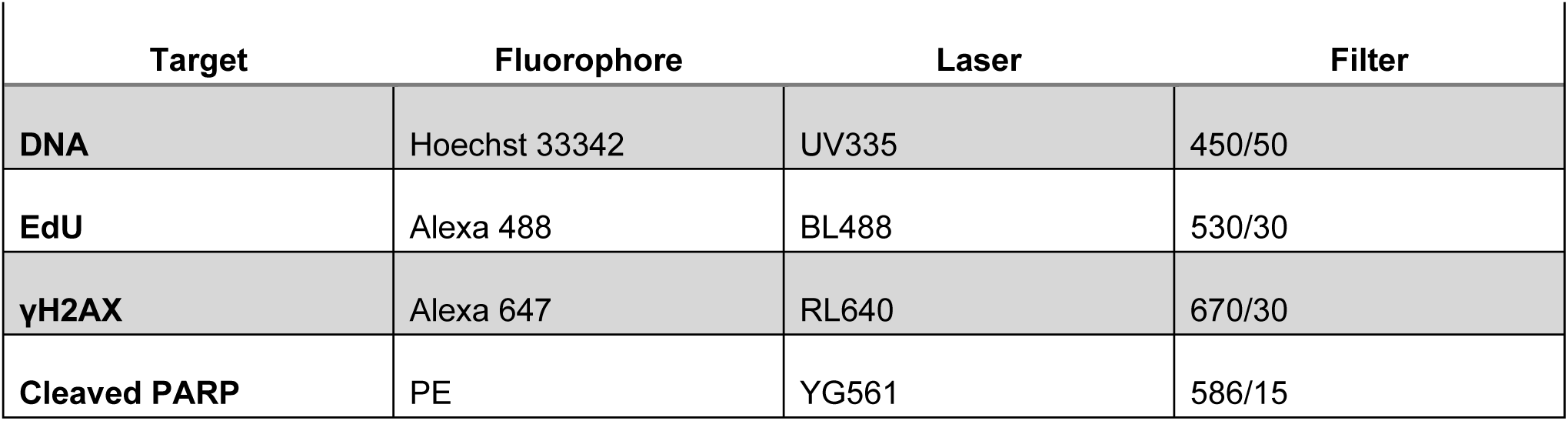
Flow Cytometry Panel Design with Lasers and Corresponding Filters.

| Target | Fluorophore | Laser | Filter |
| --- | --- | --- | --- |
| **DNA** | Hoechst 33342 | UV335 | 450/50 |
| **EdU** | Alexa 488 | BL488 | 530/30 |
| **γH2AX** | Alexa 647 | RL640 | 670/30 |
| **Cleaved PARP** | PE | YG561 | 586/15 |

Samples were analyzed in a 96-well microplate with V-shaped bottoms using the HTS in standard throughput mode. Detailed settings are provided in Table 5. Controls, including single stainings and unstained samples, were measured in tubes to allow for adjustments of PMT voltage settings, if necessary.

**Table 5.** High-Throughput Sampler (HTS) Settings for 96-Well Microplate Analysis.

| Parameter | Setting |
| --- | --- |
| **Sample Flow Rate [μl/sec]** | 0.5 |
| **Sample Volume [μl]** | 180 |
| **Mixing Volume [μl]** | 100 |
| **Mixing Speed [μl/sec]** | 180 |
| **Number of Mixes** | 2 |
| **Wash Volume [μl]** | 200 |
| **BLR Period** | 5 |

### Gating

Flow cytometry data were gated in R (v 4.5.2) using the packages flowStats (v4.22.0), flowCore (v2.22.0)^44^, flowViz (v1.74.0), gridExtra (v2.3), and tidyverse (v2.0.0). Fluorescence intensities for cleaved PARP, EdU, and γH2AX were transformed using a logical transformation to appropriately scale low and high signal distributions. Debris was excluded based on the forward and side scatter area (FSC-A and SSC-A). Single cells were identified based on the Hoechst area versus height (Hoechst-A vs Hoechst-H) of the untreated HAP1 wildtype samples. Gating was intentionally permissive to retain the apoptotic populations relevant to the study. To identify single cells with DNA damage or apoptosis, the intensity of γH2AX was plotted on the Y-axis against the intensity of cleaved PARP on the X-axis. The γH2AX threshold was defined based on the double-negative population in untreated wildtype cells and applied uniformly across all samples. The cleaved PARP threshold was adapted for each replicate, based on the double-negative distribution of the untreated wildtype samples. Cells were classified into four populations: double-negative (γH2AX−/PARP−), only-γH2AX-positive (γH2AX+/PARP−), only-PARP-positive (γH2AX−/PARP+), and γH2AX-PARP-positive (γH2AX+/PARP+).

To define the cell cycle phases, the single cells were plotted with EdU intensity on the X-axis and Hoechst intensity on the Y-axis. Cells with a high EdU intensity were defined as S phase cells, where the EdU threshold was defined based on the wildtype sample. A low EdU intensity and a low Hoechst intensity were used to define G1 phase-like cells. Approximately a two-fold higher Hoechst intensity relative to the G1-like cells and a low EdU intensity were used to define G2 phase-like cells. The Hoechst threshold that separates G1 and G2 phase-like cells changed for each sample based on the distribution of the Hoechst signal for the EdU-positive cells. The G1/G2 threshold was defined as the midpoint between the 5th and 95th percentiles of the Hoechst distribution within the EdU-positive cells. The threshold for high EdU intensities was based on the wildtype untreated cells and remained constant across all samples. Samples exhibiting technical artifacts (e.g., abnormal scatter profiles or staining inconsistencies) were excluded prior to analysis. The proportion of cells in each gate relative to the total single cells was quantified and used for downstream analyses. All gating strategies (Fig. S2d) were visually validated using scatter plots for each individual sample to ensure consistent population identification.

### Immunofluorescence microscopy

To complement the live cell growth screen and measure the effects on cell number, DNA was stained and cell numbers determined by counting the nuclei using an Immunofluorescence (IF) protocol.

### Preparation of cells and staining

The cells were harvested, counted, seeded and treated as described for the Live cell growth screen, but this time in Poly-D-Lysine coated, 96-well black, clear-bottom tissue culture-treated plates (PerkinElmer, Waltham, MA, USA). Unless otherwise noted, all following steps were performed at RT on a shaker at low speed. 48h after treatment, the cells were washed three times with 100 µl PBS and fixed by adding 50µl of 4% paraformaldehyde (PFA, Sigma-Aldrich, Merck, Darmstadt, Germany, Cat.# 158127-500G) in PBS for 15 min. Then, the fixative was removed and the reaction was quenched with 100µl of IF Quenching Buffer for 5min. The cells were then washed once with PBS and permeabilized with 0.5 % Triton X-100 in PBS (Sigma-Aldrich, Merck, Darmstadt, Germany, Cat.# T8787-250ML) for 15 min. Afterwards, the cells were washed three times for 5min each in PBS. Blocking was carried out for 1h in 100µl PBS + 3% BSA. The cells were then incubated in 50 µl PBS + 3 %BSA overnight at 4°C. The next day, the cells were washed once with 0.25% Triton X-100 in PBS for 5min, followed by three 5 min washing steps with 100 µl PBS + 3% BSA. DAPI (Invitrogen, Thermo Fisher Scientific, Waltham, USA, Cat.# D3571) staining was performed by incubating the cells with DAPI diluted in 50µl PBS + 3% BSA. Finally, cells were washed three times with PBS for 5min each and stored in 200µl PBS.

### Microscopy setup and analysis

Imaging was carried out using the Opera Phenix™ High Content Screening Microscope System (Revvity). Images were acquired using a spinning disc confocal setting with a 50 um pinhole size and 550 um pinhole spacing. Imaging was performed using the 63x water, NA 1.15 objective, resulting in a field of view of 205 * 205 um and pixel size of 200 nm in xy with an image format of 1080 * 1080 pixels. DAPI fluorescence was excited with 405 nm (30 %) and emission in the wavelength range of 435-550 nm was detected over an exposure time of 100 ms on an sCMOS camera. For each well, 13 fields were acquired with a stack of 5 planes and a distance of 0.5 µm between each plane.

Analysis of the microscopy data was performed in the Harmony software (v.5.2). The image stacks were combined, using maximum projection and basic flatfield correction. Nuclei were identified based on the DAPI channel, using method M. The expected diameter of nuclei was set to 15 µm, the splitting sensitivity to 0.13 and the common threshold to 0.2. From this population, all objects touching the border were removed. To remove DAPI positive objects that are no real nuclei, all objects with a roundness under 0.72, an area smaller than 50 µm^2^, and an area larger than 690 µm^2^ were excluded. This new population was called ‘All Nuclei’. An additional filtering step was used, excluding all nuclei with an area over 300 µm^2^ to make sure all nuclei in this dataset are single nuclei and not multiple nuclei stuck together. This population was called ‘Single Nuclei’. The average nucleus area and total nuclei count were directly obtained from the Harmony software based on the segmented ‘Single Nuclei’ population.

Analysis of the raw data were performed in Python (v3.13.12) using the NumPy (v2.3.4)^45^, pandas (v2.3.3), SciPy (v1.16.3)^46^, statsmodels (v0.14.5), matplotlib (v3.10.7)^47^, and seaborn (v0.13.2)^48^ packages. A readout was only considered a valid value, if at least 20 cells were measured in that well. Bar plots were created using the raw values and the significance was calculated using the student’s t-test comparing the knockout to wildtype, where multiple testing correction for FDR was applied.

### Transcriptome analysis

#### Preparation of samples and sequencing

The indicated cell lines were seeded, treated and harvested as described for the flow cytometry screen, with the difference that the cell culture media was discarded during harvesting. After cell harvesting, the cell pellet was washed once with DPBS and stored at -80°C, until further use. Subsequently, total RNA was isolated using the RNeasy Mini Kit (Qiagen, Hilden, Germany), according to the manufacturer’s protocol. After an initial quantification of the RNA levels with the Spectrophotometer / Fluorometer DS11 FX+ (DeNovix, Wilmington, USA), total RNA samples were submitted to the IMB Genomics Core Facility for library preparation and sequencing. NGS library prep was performed with Illumina’s Stranded mRNA Prep Ligation Kit following Stranded mRNA Prep Ligation ReferenceGuide (October 2023) (Document 1000000124518 v04). Libraries were prepared in 3 equal batches with a starting amount of 1000 ng and amplified in 8 PCR cycles. Two post PCR purification steps were performed to exclude residual primer and adapter dimers. Libraries were profiled in a D5000 ScreenTape Assay on a TapeStation 4200 (Agilent technologies) and quantified using the Qubit 1x dsDNA HS Assay Kit, in a QubitFlex Fluorometer (Invitrogen by Thermo Fisher Scientific). Library preparation of sample 68 was repeated in batch 3 with a different UDI. All 136 libraries were pooled in equimolar ratio aiming for at least 25M reads per sample and sequenced on 2 NextSeq 2000 P4 (100 cycles) flow cells, SR for 1x 116 cycles plus 2×10 cycles for the index reads and 1 dark cycle upfront R1.

#### Processing of reads and read count generation

Raw data demultiplexing has been performed using Illumina’s bcl-convert software v.4.1.5 and overall sequence quality and rRNA content is assessed with FastQC v0.11.9^49^ and FastQ Screen v0.15.2^50^. Reads were trimmed for Illumina adapter sequences using cutadapt v.4.0^51^ and subsequently mapped to Gencode reference genome GRCh38.p14 (release 45) with STAR v2.7^52^ with default parameters (except --outFilterMismatchNoverLmax 0.04 and --outFilterMismatchNmax 999). Only uniquely mapped reads were used for downstream analysis. Coverage signal tracks (bigWigs) of primary alignments were generated using DeepTools v3.5.1^53^. Read summarization at gene level (exon reads counted) is performed using Subread featureCounts v.2.0^54^ with default parameters.

#### Differential gene expression analysis, clustering, and GO enrichment analysis

The differential gene expression analysis was performed on the raw gene counts using the DESeq v1.50.2^55^ R package using the design formula ’∼ genotype + treatment + genotype:treatment’, from which the ‘genotype:treatment’ parameter modeled the interaction effect of the change in gene expression upon compound treatment for a given BAF subunit knockout compared to DMSO and the WT cell line. P-values were adjusted for multiple testing for each comparison using the Benjamini-Hochberg procedure. Using the clusterProfiler v4.18.1^56^ package to extract the gene type information, the expression counts of all tested protein-coding genes, that are significantly differentially expressed in at least one condition (KO cell line + compound) with an adjusted p-value < 0.001 were plotted as a heatmap using the pheatmap v1.0.13 with k-means clustering method applied across the genes. The Gene Ontology term enrichment analysis was then performed on the DEGs within each cluster using the enrichGO function of the clusterProfiler package, applying a significance threshold of p-value < 0.05 after adjusting for FDR. Gene lists from cellular pathways pertaining to DNA damage repair, cellular growth signaling and cell cycle checkpoints were created by downloading their respective protein names and UniProt IDs from the Reactome pathway browser release 95 accessed November 11, 2025^57^. The DEGs were then subsetted using these gene lists and filtered for DEGs significant in at least one condition with an adjusted p-value < 0.01. These significant hits were visualized in a heatmap using the pheatmap R package with k-means clustering for each pathway.

### Analysis of external datasets

#### Olivieri dataset

Data of 31 genome-wide CRISPR-Cas9 screens against 27 genotoxic agents in the hTERT-immortalized RPE1 cell line, where the amount of sgDNA per sample indicated sensitivity, was downloaded from Oliveri et al.^28^ To identify hits in the screen, the z-score cutoff of -3 and 6 were applied to define sensitizing hits and resisting hits, respectively, along with a false discovery rate (FDR) < 15%. The data was filtered to only include the following genes encoding BAF complex subunits: ARID1A, ARID1B, ARID2, BRD7, PBRM1, PHF10, BRD9, SMARCA4, SMARCC1, SMARCD1.

#### PRISM dataset

Sensitivity measurements of cancer cell lines from 24 tumor types treated primarily with anti-cancer drugs and compounds in clinical trials were obtained from the PRISM repurposing dataset^27^. In the PRISM assay, individual cancer cell lines are uniquely barcoded and pooled prior to treatment with chemical perturbagens. Following drug treatment for five days, the relative abundance of each cell line is quantified by measuring barcode-associated fluorescence signals, where barcode abundance reflects cell viability. Median Fluorescence Intensity (MFI) values therefore represent the relative abundance of each cell line after treatment and were used as a proxy for drug sensitivity. The dataset was generated in two screens: a primary screen where 919 cell lines were treated with a single concentration of each compound (6,658 compounds total), followed by a secondary screen where 1,448 compounds with activity in the primary screen were retested at eight doses. In the primary screen, MFI values at the single drug concentration were used to assess drug sensitivity, whereas in the secondary screen MFI measurements across eight doses were used to derive dose–response relationships.

Cell line annotations and expression data were obtained from the DepMap CCLE dataset of the 22Q2 release (Accessed 17-04-2024). It contained expression level scores for 16,383 protein-coding genes of 1,405 cancer cell lines. Values are inferred from RNA-seq data and are reported after log2 transformation using a pseudo-count of 1. The PRISM dataset was filtered for BAF subunit genes, selecting only the top and bottom 20th percentile of cell lines in terms of expression levels for each BAF subunit, defining two groups: the high expression group and the low expression group of cell lines for each BAF subunit. The two groups of cell lines were compared to each other with respect to the MFI values for a given treatment. Given the non-normal distribution of drug response values across the cell lines, differences in MFI values between both groups were assessed using the Mann– Whitney U-test. The Mann–Whitney U-test was performed using the mannwhitneyu function from the stats module from the scipy (v1.16.3) package and the mean difference of the two groups (low _-_high expression) was calculated as an effect size, followed by Benjamini-Hochberg correction of the results. Significant results were identified as having an adjusted p-value < 0.01 in both the primary and secondary screen. If the low expression group had a lower MFI than the high expression group, we assumed sensitivity of cells with respect to a given BAF subunit, and vice versa for resistance. The mean difference for each treatment and BAF subunit combination was visualized as a heatmap (seaborn (v0.13.2), matplotlib (v3.10.7) packages) with positive values indicating resistance and negative values indicating sensitivity.

#### HAP1 KO RNA-seq dataset

The gene-level expression count data for an array of HAP1 BAF KOs was downloaded from https://medical-epigenomics.org/papers/schick2018/#d^8^ and analyzed for differential gene expression using two different approaches, both performed in R (v 4.5.2). In the first approach, the expression of a given gene for a given BAF subunit KO was compared to its expression in all other BAF KOs and WT, to quantify to which extent this gene is preferentially up- or down-regulated in the respective BAF KO in comparison to all other BAF KOs. Quantification was done by applying a pseudo count of 1 and log2 transformation on the count data for protein-coding genes. Then the Preferential Expression Score (PES) for a given gene and BAF KO was calculated as follows^24^

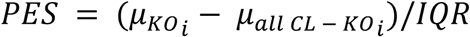

where 𝜇_𝐾𝑂𝑖_ is the mean expression count for a given gene in a given BAF subunit KO cell line, 𝜇_𝑎𝑙𝑙 𝐶𝐿 − 𝐾𝑂𝑖_ represents the mean of expression of that gene in all other cell lines including wildtype, and 𝐼𝑄𝑅 is the interquartile range of the expression of that gene in all cell lines. Genes with |PES| > 1 in at least one cell line were visualized in a heatmap with k-means clustering using the ComplexHeatmap (v2.26.1)^58^ R package. GO term enrichment analysis was performed for each cluster using the enrichGO function from the clusterProfiler (v4.18.3) package. Heatmap annotations were manually curated based on significantly enriched GO terms, with an adjusted p-value < 0.05 after adjusting for FDR.

In the second approach, the differential gene expression analysis was performed using the raw gene counts as input for the DESeq2 (v1.50.2) R package, which modeled the change in gene expression upon a given BAF subunit KO compared to the wildtype cell line. P-values were corrected for multiple testing for each individual comparison using the Benjamini-Hochberg procedure. The results were filtered for protein-coding genes with an adjusted p-value < 0.01 and log2 fold-change > 1 or < -1 in at least one KO. These were visualized in a heatmap using the R package pheatmap (v1.0.13) with k-means clustering applied across the genes. The GO term enrichment analysis was then performed on the genes within each cluster using the R package clusterProfiler (v4.18.1), where a significance threshold of p-value < 0.05 was applied after adjusting for FDR. The number of up- and downregulated differentially expressed protein-coding genes in each KO were summarized in a bar plot using the ggplot2 (v4.0.1)^59^ package. Using the Reactome pathway-specific gene lists, as previously described for the analysis of the compound treated transcriptome data, the original differentially expressed genes were subsetted for each pathway and filtered for significantly differentially expressed genes (adjusted p-value < 0.05) in at least one KO. These were then visualized in a heatmap with k-means clustering applied across the genes for each pathway. The number of significantly up- and downregulated differentially expressed genes (adjusted p-value < 0.05 and log2FC > 0 or < 0) in each knockout for each pathway were summarized in a bar plot.

### Incucyte data analysis

#### Analysis of data from untreated samples

The untreated confluency data was fitted to a Non-Linear Mixed Effects Model (NLMM) using the nlme (v3.1-168)^60–62^ R package from which the mean estimated relative growth rate, initial confluency, and maximum capacity were extracted per condition. The selfStart function SSlogis from the stats (v4.5.2) package was re-parametrized to create the following logistic growth model:

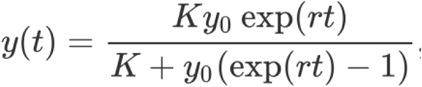

The parameters Asym, xmid, and scal from the SSlogis function were transformed into the parameters K (maximum capacity), y0 (initial confluency at time 0 hr) and r (relative growth rate) of the logistic growth function, respectively, in order to extract the coefficients along with their standard errors directly. The random effects accounted for the covariance caused by the biological replicates avoiding pseudo replication. To compare the KO to wildtype, the model fitted both conditions together, where the genotype was defined as a fixed effect. From this the significance of the genotype effect could be extracted (H0: log2FC(K) = 0, log2FC(r) = 0). We corrected for multiple testing using the Benjamini-Hochberg correction for each coefficient. The log2 fold change of the KO versus WT for each model estimate was visualized using the matplotlib (v3.10.7) package in Python (v3.13.12).

#### Analysis of data from treated samples

The following phenotypes were extracted from the confluency data, which use the amount of time that has passed since the treatment was administered, including: the confluency after 24hr, 48hr, 72hr, and 96hr of treatment as well as the ratio of the confluency after 96hr over that after 48hr of treatment, in order to capture the change in growth behavior at the later timepoints. These were then normalized to the DMSO-treated sample from the same biological replicate per cell line, in order to account for batch effects. By linearly interpolating between the tested concentrations and applying an adaptive quadrature approach, the Area Under the Curve (AUC) of the dose-response curve for each compound and biological replicate was then calculated per cell line. Using the limma (v3.66.0)^63^ R package, the log2 fold-change was calculated and an empirical Bayes moderated paired t-test was performed on the AUC values comparing KO to WT. The resulting p-values were corrected for multiple testing using the Benjamini-Hochberg procedure. The unsupervised clustering method k-means was applied on these log2 fold-change values across all 5 phenotypes and visualized as a heatmap using the ComplexHeatmap (v2.26.0) package.

### Flow cytometry data analysis

Downstream analyses were performed in Python (v3.13.12) using the NumPy (v2.3.4), pandas (v2.3.3), SciPy (v1.16.3), statsmodels (v0.14.5), matplotlib (v3.10.7), and seaborn (v0.13.2) packages.

### Analysis of cell cycle data

Percentages of cells in a given cell cycle phase from the compound-treated samples were normalized to the respective DMSO-treated samples from the same HAP1 BAF KO cell line and biological replicate and converted to log2 scale. Statistical comparisons between knockout and wildtype samples were performed across biological replicates using an unpaired two-sided Student’s t-test on the log2-transformed values, followed by FDR correction. These were visualized as heatmaps. Bar plots display the average percentages across four biological replicates per condition without normalization to DMSO.

### Analysis of apoptosis and DNA damage data

As there was a small amount of apoptotic and DNA damage positive cells in the untreated conditions, normalization was performed by subtracting the corresponding DMSO-treated population from the compound-treated samples from the same cell line. Statistical comparisons between KO and wildtype samples were performed using an unpaired two-sided Student’s t-test on the normalized values, followed by FDR correction. Heatmaps display the mean difference between KO and wildtype for the normalized values across the four biological replicates. Bar plots display the non-normalized percentages, averaged across the four biological replicates per condition. Here the statistical significance comparing KO to wildtype was calculated on the non-normalized values using an unpaired Student’s t-test followed by FDR correction. More detailed analyses linking to which cell cycle phase the PARP-positive and γH2AX-positive cells were in, were done as follows. PARP-positive and γH2AX-positive populations were defined as 100%, and the distribution of cell cycle phases within these populations was quantified. Stacked bar plots were generated to compare the mean cell cycle distribution of PARP- or γH2AX-positive populations to their respective negative populations in DMSO-treated samples per condition across the four biological replicates.

## Supplementary Figures

**Figure S1:**
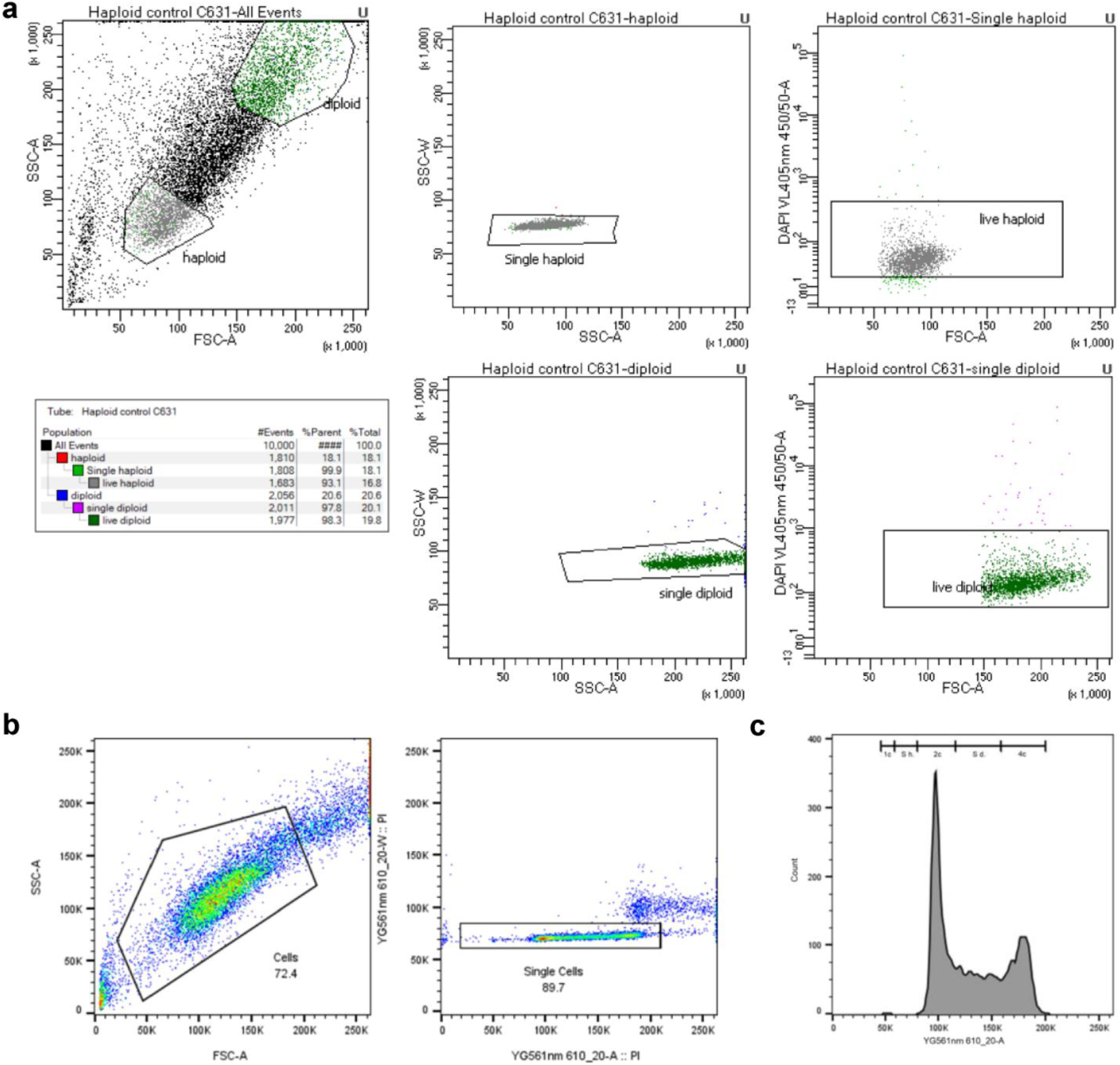
Isolation and validation of diploid HAP1 cell lines. **a.** FACS gating strategy used for sorting live single haploid (control HAP1 WT) or diploid cells, showing size/granularity (FSC-A vs SSC-A), haploid/diploid separation, and exclusion of DAPI-positive dead cells. **b.** FACS gating strategy used for downstream analysis of fixed diploid HAP1 cells. **c.** Propidium iodide (PI) DNA content histogram of the sample shown in **b.,** illustrating cell cycle distribution and confirming diploid enrichment. 1C, 2C, 4C indicate relative DNA content; S h. = S phase haploid cells, S d. = S phase diploid cells

**Figure S2:**
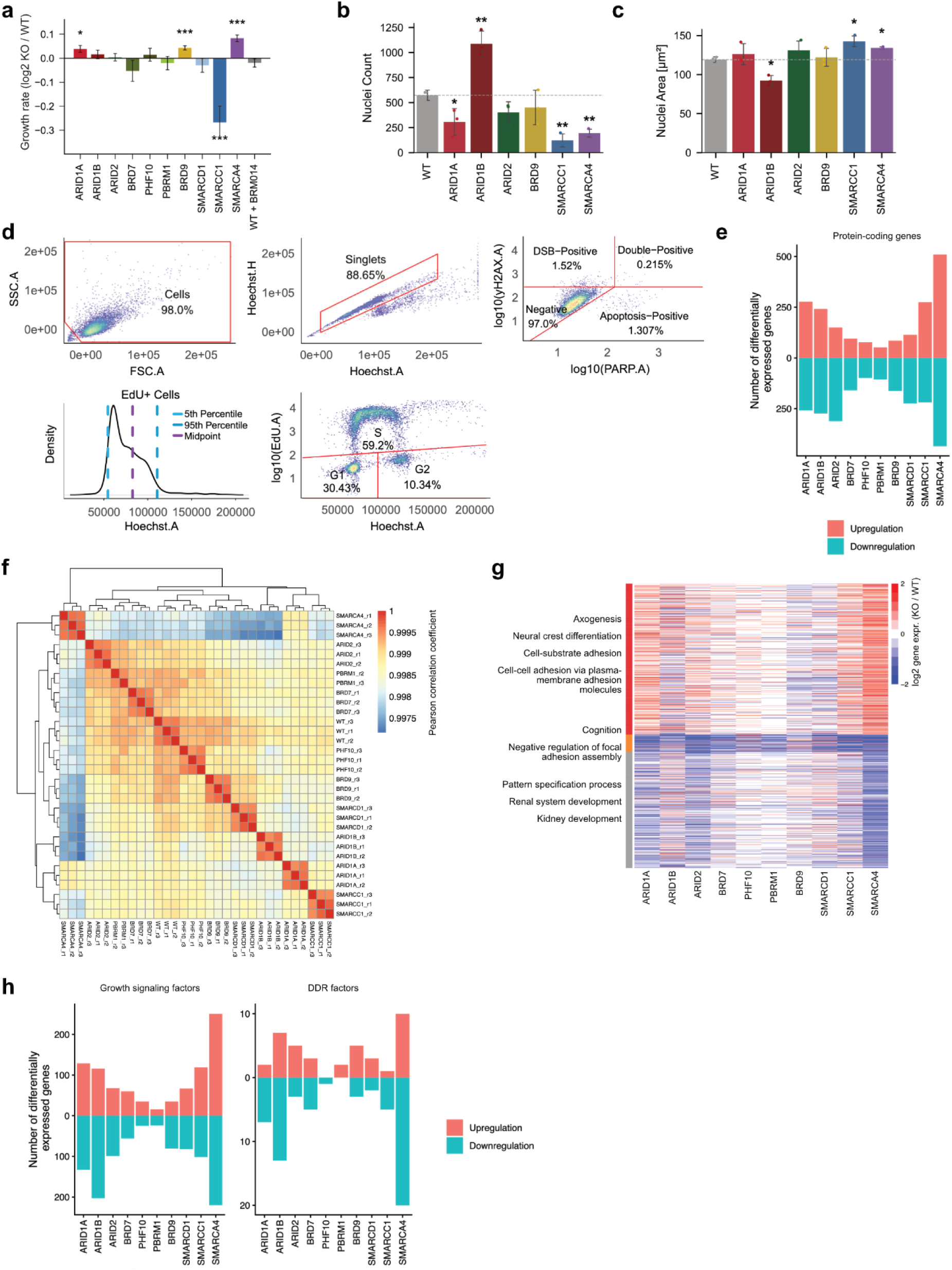
Cellular phenotypes and transcriptomic differences across HAP1 BAF KO lines. **a.** Bar plot showing estimated mean max. capacity from fitted confluency curves of BAF KO cell lines relative to WT. Error bars indicate STE, n=3. **b.** Bar plot showing mean nuclei count per well for WT and BAF KO cells obtained by immunofluorescence imaging. Dashed line indicates WT. Error bars show STE, n=3. Significance was determined using t-test, p-values corrected for multiple testing, * p-value < 0.05, ** p-value < 0.01, *** p-value < 0.001. **c.** Mean of nuclei area measured by immunofluorescence imaging for WT and BAF KO cells using DAPI stain. Dashed line indicates WT. Error bars show STE, n=3. Statistics as in **b**. **d.** Schematic illustrating flow cytometry gating strategy used for analysis of cell cycle, apoptosis, and DNA damage. Cells were first gated based on forward and side scatter (FSC-A vs SSC-A), followed by singlet selection using Hoechst height and area. DSB positive and apoptotic cells were quantified using γH2AX and cleaved PARP (PARP.A) staining, respectively. Quadrants indicate γH2AX-positive (DSB-positive), cleaved PARP-positive (apoptotic), γH2AX-and cleaved PARP-positive (double-positive), and γH2AX- and cleaved PARP-negative (negative) cell populations. Cell cycle phase distribution was determined using Hoechst DNA content together with EdU incorporation to identify G1, S, and G2 populations as indicated. **e.** Bar plot showing the number of differentially expressed protein-coding genes identified by RNA-seq for each BAF KO cell line compared to WT. **f.** Pearson correlation heatmap of transcriptome profiles across WT and BAF KO cell lines based on RNA-seq data. Hierarchical clustering reveals similarities and differences between transcriptional profiles of individual KO lines. **g.** Heatmap clustered by differential expression of protein-coding genes across HAP1 BAF KO panel compared to WT. Selected enriched GO biological process categories for the three clusters are indicated. **h.** Bar plots showing the number of differentially expressed genes involved in growth signaling pathways (left) and DNA damage repair (DDR) pathways (right) across BAF KO cell lines compared to WT.

**Figure S3:**
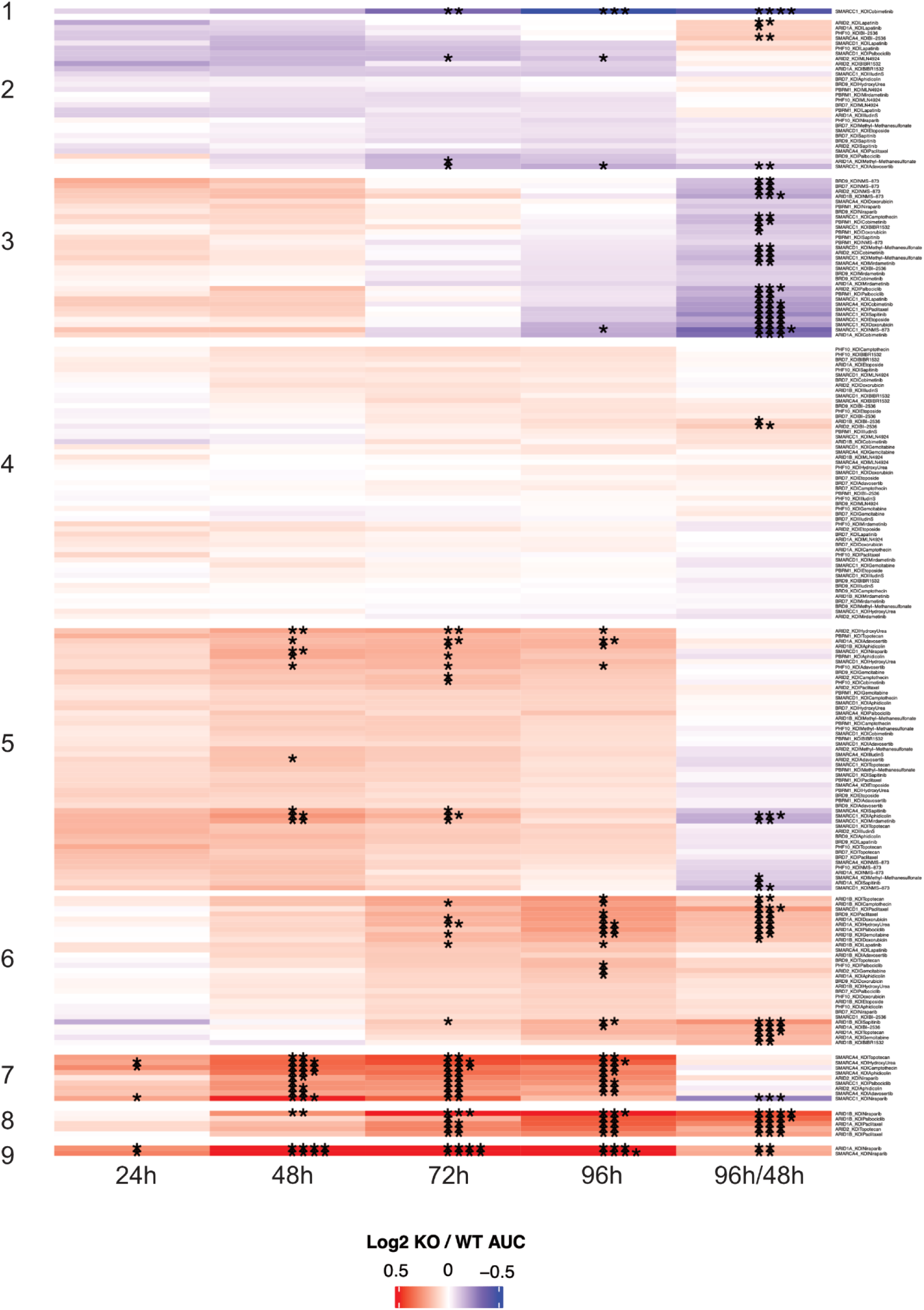
Heatmap showing Area Under the dose response Curve (AUC) of confluencies measured for BAF KO over WT for different treatments and time points after treatment, normalized each to DMSO and clustered into nine groups. Confluency measurements were obtained from live-cell imaging microscopy. Conditions were clustered using k-means across the five time-dependent readouts. Significance determined using t-test, p-values corrected for multiple testing, * p-value < 0.1, ** p-value < 0.05, *** p-value < 0.01 and **** p-value < 0.001, n=3.

**Figure S4:**
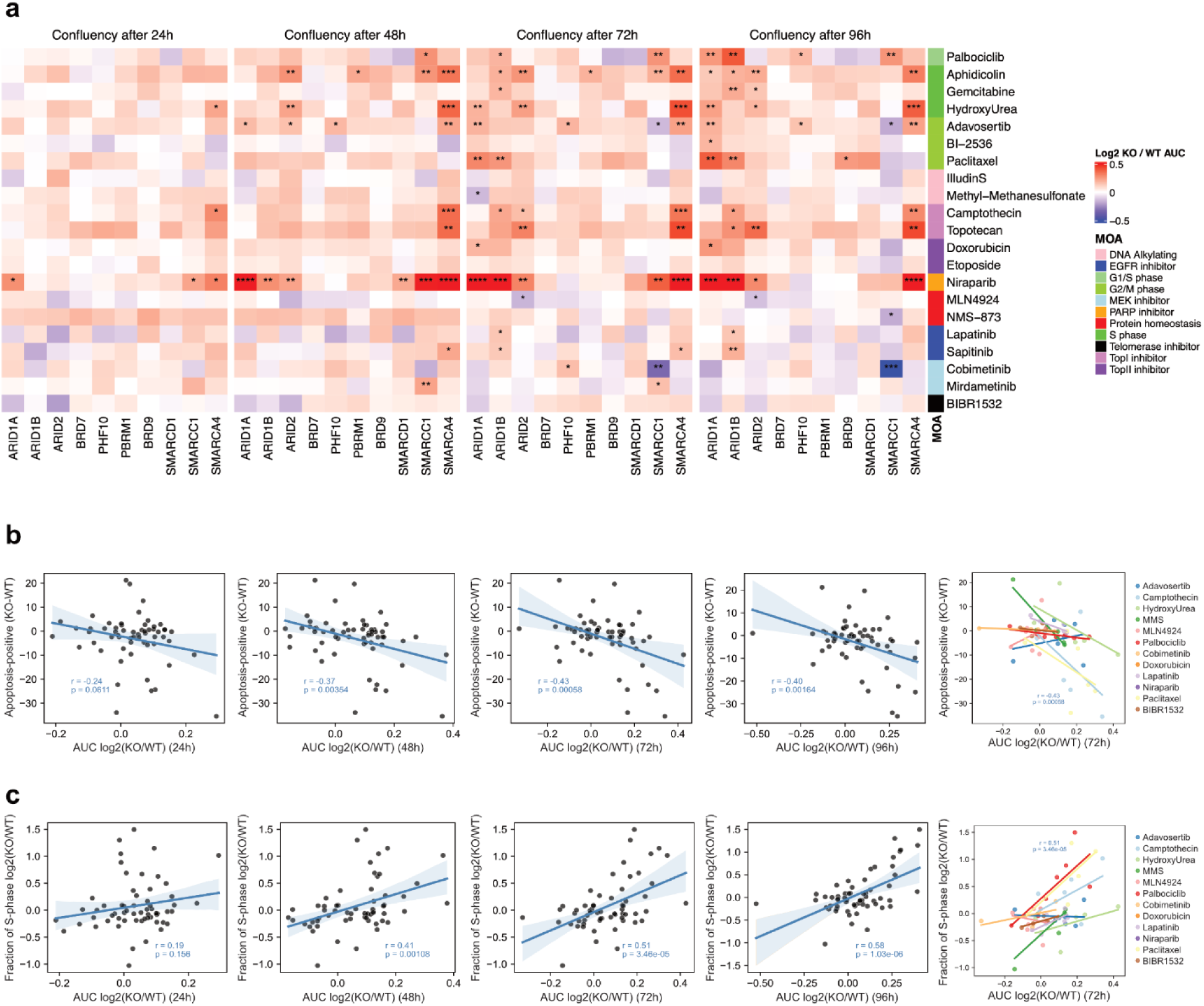
a. Heatmaps show mean log2 fold-change of Area Under the dose response Curves (AUC) of confluency across treatments and the HAP1 BAF KO panel compared to treated WT after 24h, 48h, 72h and 96h of treatment (from left to right), each compared to treatment with DMSO. Confluency measurements were obtained from live-cell imaging microscopy. Significance determined using moderated paired t-test, p-values corrected for multiple testing, * p-value < 0.1, ** p-value < 0.05, *** p-value < 0.01, **** p-value < 0.001, n=3. **b.** Scatterplots of fraction of cleaved PARP-positive BAF KO cells minus WT (Flow cytometry data) and relative AUC KO/WT for different time points after treatment (Incucyte data) as indicated. **c.** Scatterplots of fraction cells in S phase in treated BAF KO cells compared to treated WT (each compared to DMSO) (Flow cytometry data) and relative AUC KO/WT for different time points after treatment (Incucyte data) as indicated.

**Figure S5:**
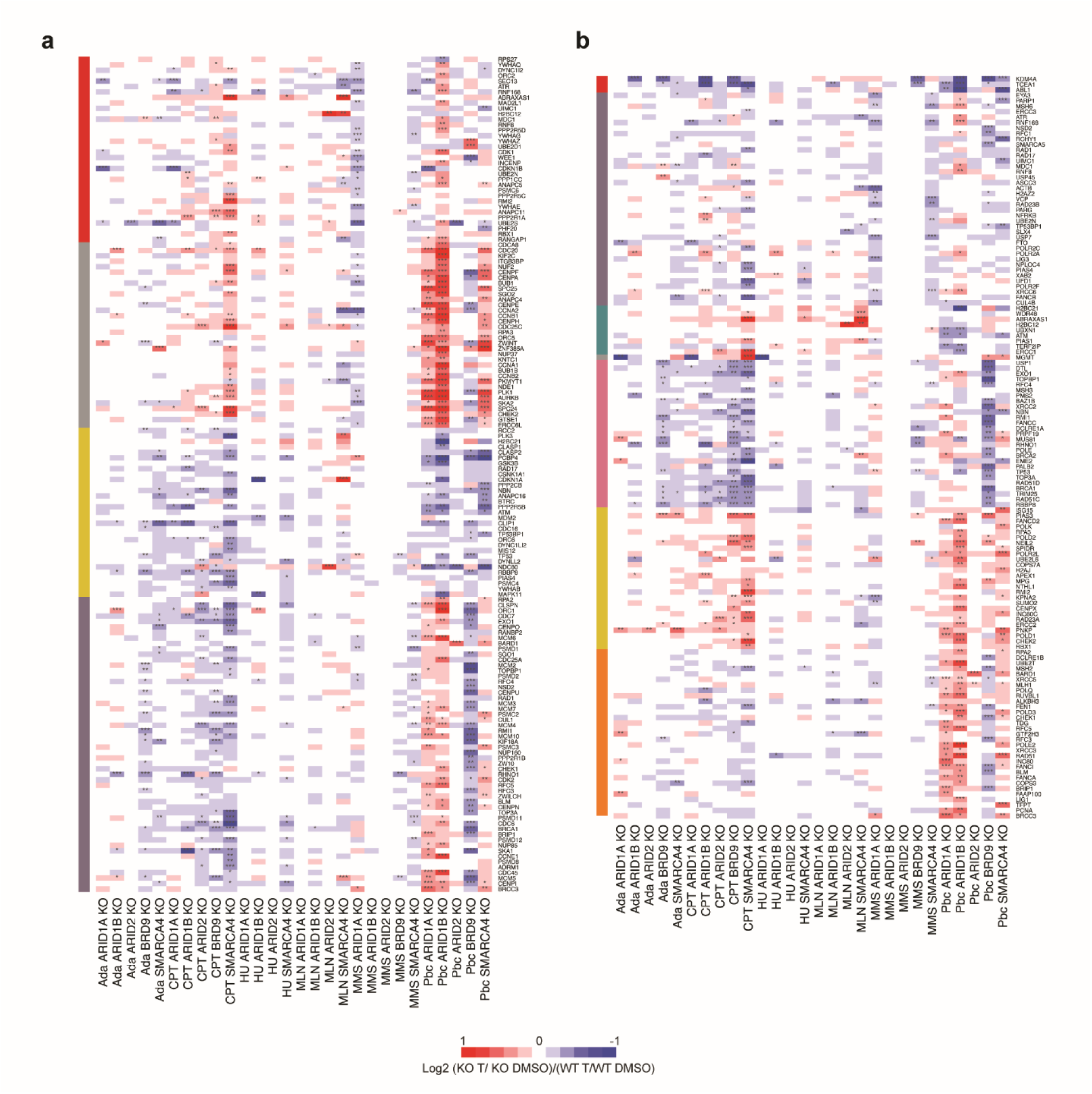
Differential expression of genes upon BAF KO and treatment in comparison to WT treated and both compared to treatment with DMSO for **a.** Reactome pathway cell cycle checkpoint genes and **b.** Reactome DDR pathway genes. Genes significant for at least one BAF KO and treatment are shown based on an adjusted p-value < 0.01. K-means clustering was applied and significance determined using DESeq2, p-values corrected for multiple testing, * p-value < 0.05, ** p-value < 0.01, *** p-value < 0.001, n=3.

